# Lymphocyte networks are dynamic cellular communities in the immunoregulatory landscape of lung adenocarcinoma

**DOI:** 10.1101/2022.08.11.503237

**Authors:** Giorgio Gaglia, Megan L. Burger, Cecily C. Ritch, Danae Rammos, Yang Dai, Grace E. Crossland, Sara Z. Tavana, Simon Warchol, Alex M. Jaeger, Santiago Naranjo, Shannon Coy, Ajit J. Nirmal, Robert Krueger, Jia-Ren Lin, Hanspeter Pfister, Peter K Sorger, Tyler Jacks, Sandro Santagata

**Author notes:** These authors contributed equally to this work (Giorgio Gaglia, Megan L. Burger). Equal contribution (Peter K Sorger, Tyler Jacks, Sandro Santagata). Lead contact: Sandro Santagata, Brigham and Women’s Hospital, 60 Fenwood Road, Boston, Massachusetts 02115.

## Abstract

Lymphocytes play a key role in immune surveillance of tumors, but our understanding of the spatial organization and physical interactions that facilitate lymphocyte anti-cancer functions is limited. Here, we used multiplexed imaging, quantitative spatial analysis, and machine learning to create high-definition maps of tumor-bearing lung tissues from a Kras/p53 (KP) mouse model and human resections. Networks of directly interacting lymphocytes (‘lymphonets’) emerge as a distinctive feature of the anti-cancer immune response. Lymphonets nucleate from small T-cell clusters and incorporate B cells with increasing size. CXCR3-mediated trafficking modulates lymphonet size and number, but neoantigen expression directs intratumoral localization. Lymphonets preferentially harbor TCF1+/PD1+ progenitor CD8 T cells involved in responses to immune checkpoint blockade (ICB). Upon treatment of mice with ICB therapy or a neoantigen-targeted vaccine, lymphonets retain progenitor and gain cytotoxic CD8 T-cell populations, likely via progenitor differentiation. These data show that lymphonets create a spatial environment supportive of CD8 T-cell anti-tumor responses.

## INTRODUCTION

During cancer progression, immune cells proliferate, adapt, and migrate in an attempt to impede tumor spread (Hanahan, 2022; Nirmal et al., 2022). Tumor cells respond by inducing programs that suppress immune cell function and migration (Bailey et al., 2021). Detailed characterization of the functional states of immune cells and their spatial organization relative to tumor cells is needed to identify the features of anti-tumor immunity (Pelka et al., 2021). One way to accomplish this is using the analytical methods and computational approaches that constitute highly multiplexed spatial profiling, an emerging field that seeks to provide quantitative descriptions of i) the identities and molecular characteristics of immune, tumor and stromal cells, ii) the physical and chemical factors that influence the spatial organization of these cell types, and iii) how spatial features change over time and space and in response to therapy (Baertsch et al., 2022; Bodenmiller, 2016; Lewis et al., 2021).

Genetically engineered mouse models (GEMMs) of cancer represent an important tool for studying the effects of genetic and chemical perturbations on tumors to better understand mechanisms of oncogenesis and therapy (DuPage and Jacks, 2013; Yap et al., 2021). Tumors in GEMMs are initiated de novo from single transformed cells, and their development recapitulates many histological and molecular features of human cancer (Kersten et al., 2017). Importantly, the kinetics of tumor growth in immunocompetent GEMMs allows for interactions between tumor and immune cells that are more relevant to human disease than transplant cancer models. GEMMs can also be sampled longitudinally over the course of tumor progression and following treatment with immunotherapies (Yap et al., 2021). By contrast, human tumors are typically studied at intermittent time points dictated by clinical necessity.

The Kras/p53-mutant (KP) model of lung adenocarcinoma, which includes several variants, is prototypical of GEMMs having many of the features of human cancer. In the KP model, tumorigenesis is synchronously initiated in multiple cells by intratracheal delivery of lentiviruses containing Cre recombinase into Kras^LSL-G12D/+^;p53^fl/fl^ animals (DuPage et al., 2009; Johnson et al., 2001). This gives rise to ∼10-15 tumor nodules per 2-dimensional lung cross-section that progress from hyperplasia to adenoma to adenocarcinoma over the course of 1-5 months. Because these tumors have a much lower rate of somatic mutations than human lung cancers, they are not highly immunogenic (McFadden et al., 2016). To overcome this limitation, CD8 T-cell neoantigens can be introduced by way of the tumor-initiating lentiviruses. In the LucOS variant of the KP model, the SIINFEKL (SIIN) epitope derived from chicken ovalbumin and the synthetic peptide SIYRYYGL (SIY) are expressed as a fusion to luciferase in tumor cells (DuPage et al., 2011). Conventional single-marker IHC analysis of tumor-bearing lung tissue from KP LucOS versus control (KP Cre) mice has shown expression of LucOS substantially increases the number of tumor-infiltrating CD8 T cells in early adenomas. However, despite the engagement of immunosurveillance mechanisms, tumor growth rebounds within weeks with a concomitant decline in the CD8 T-cell response (DuPage et al., 2011).

It is well established that dissociative single-cell methods such as single-cell RNA-sequencing (scRNA-seq), cytometry by time of flight (CYTOF), and multiparameter fluorescence activated cell sorting (FACS) can provide deep insight into tumorigenesis and immunosurveillance in GEMMs (Liu et al., 2021). However, these methods lack detailed information on cell-cell interactions, the position of immune cell populations and tumor cells relative to one another, and the roles played by tissue structures. Positional information can be obtained from conventional histology (using hematoxylin and eosin stains; H&E) and IHC but such approaches do not provide sufficient molecular information to precisely identify and phenotype cells. Deep spatial profiling has recently become possible through the use of highly-multiplexed antibody-based tissue imaging methods such as MxIF, CODEX, 4i, mIHC, MIBI, IMC, and cyclic immunofluorescence (CyCIF) (Angelo et al., 2014; Gerdes et al., 2013; Giesen et al., 2014; Goltsev et al., 2018; Gut et al., 2018; Lin et al., 2018; Tsujikawa et al., 2017).

In this study, we used multiplexed tissue profiling methods to examine the spatial features of tumor-immune interactions in KP LucOS lung tumors, including when chemokine-mediated trafficking was modulated as well as when tumors were treated with a neoantigen-targeted vaccine or immune checkpoint blockade (ICB) therapy. Intratumoral lymphocyte networks (lymphonets) were identified as key components of the neoantigen-directed immune response against early lesions. Critically, these networks harbor stem-like, progenitor CD8 T cells that have previously been associated with therapeutic response to ICB (Philip and Schietinger, 2022). Lymphonets were similarly abundant across early-stage human lung cancer resections and structurally distinct from tertiary lymphoid structures (TLS), much larger B-cell rich lymphoid structures that have been strongly associated with prognosis and response to immunotherapy (Schumacher and Thommen, 2022). These data establish generally useful methods for spatial analysis of GEMMs and identify lymphonets as components of functional T-cell responses in early tumor lesions and following immunotherapy.

## RESULTS

### Spatial analysis of the KP GEMM tumor-immune microenvironment by multimodal data integration

To generate high content spatial maps of tumor and immune cell interactions in KP lung tumors under multiple biologically informative conditions, KP mice were exposed to different tumor-initiating lentiviruses via intratracheal delivery and treated with immune therapies (**Figure 1A**). Six to nine weeks after tumor initiation, H&E staining, mRNA *in situ* hybridization (ISH) and 24-plex CyCIF (**Table S1**) (Lin et al., 2018) were performed on serial whole-slide sections (∼1 cm^2^) of formalin-fixed paraffin embedded (FFPE) tissue containing 2 or 3 lung lobes. Histopathological annotation of H&E images provided data on the position of tumor nodules and normal anatomic structures, including large and mid-sized airways as well as blood vessels (**Figure S1A**). RNA *in situ* hybridization (ISH) provided information on critical chemokines (e.g., CXCL9 and CXCL10) that are difficult to image in tissue using antibodies. For CyCIF, a 24-plex antibody panel was developed that included lineage-specific transcription factors such as Nkx2-1 (also referred to as TTF-1) and the intermediate filament protein pan-cytokeratin (Pan-CK), both of which mark epithelial/tumor cells, and vimentin (Vim), which marks all mesenchymal cells, as well as surface markers expressed on specific lymphoid and myeloid cell types (CD45, CD3e, B220, NKp46, CD11b, CD11c, Ly6G, CD103) (**Figures 1B-1D**, and **S1B**). These immune markers made it possible to delineate cell types with increasing depth, separating lymphoid and myeloid lineages, and subdividing them into T-cell, B-cell, natural killer (NK)-cell, neutrophil, dendritic cell, alveolar macrophage, and tumor-associated macrophage (TAM) populations (**Figure 1D**, see **Figure S1C** for a classification dendrogram). Additional markers (CD4, CD8, and Foxp3) made it possible to distinguish T helper (Th), T cytotoxic (Tc), and T regulatory (Treg) cell populations, and functional markers were used to define their cell state, including Ki-67 (proliferation), cytotoxicity markers granzyme B (GzmB) and perforin (Prf), inhibitory receptors PD1 and TIM3, and the T-cell transcription factor (TCF1), a key regulator of T-cell function and differentiation (**Figures 1C, 1D** and **S1C**).

**Figure 1.**
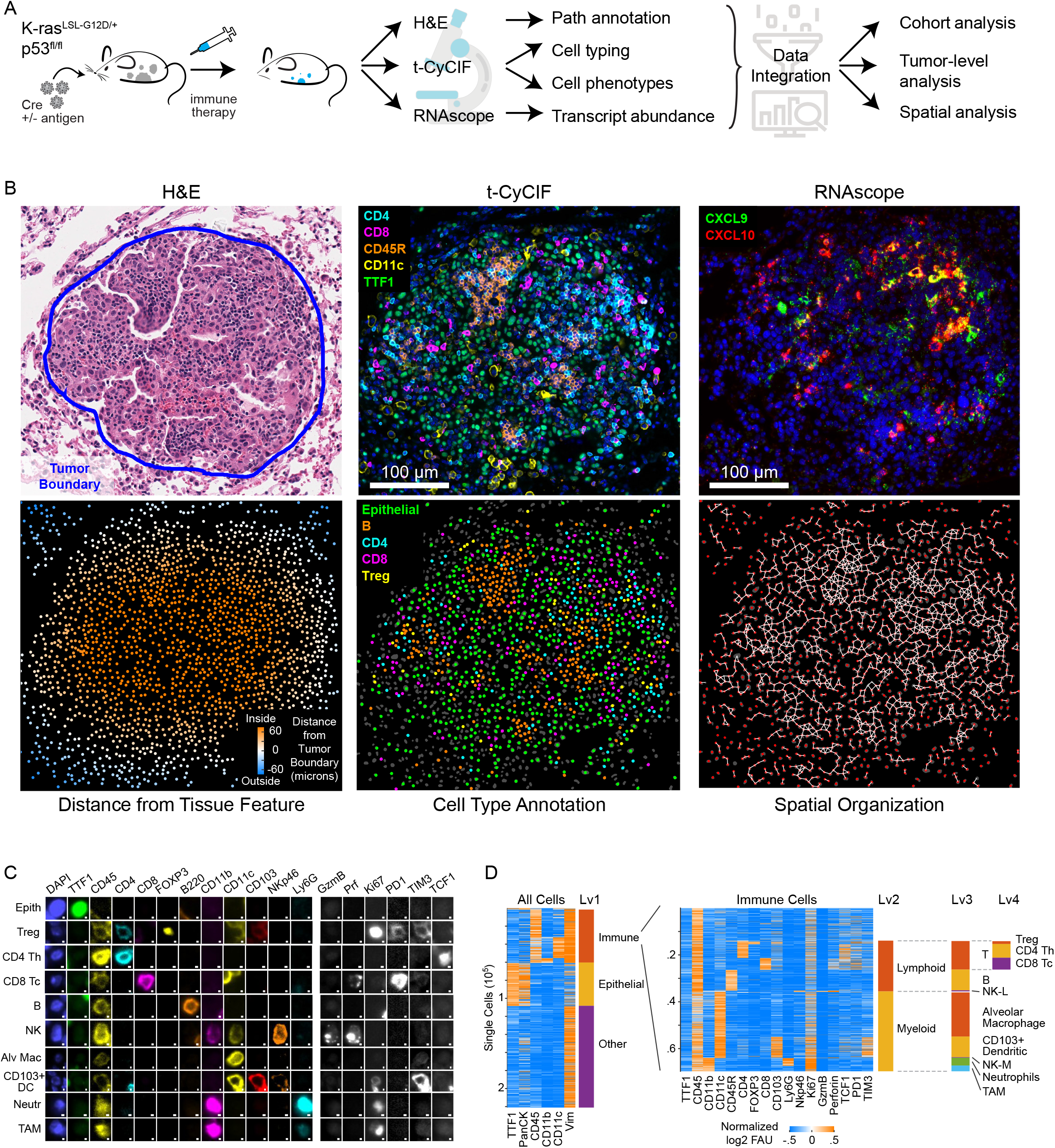
Spatial analysis of the KP GEMM tumor-immune microenvironment by multimodal data integration. (A) Schematic of the KP GEMM of lung cancer, treatments, and multi-modality data integration and analysis. (B) Representative images acquired from a tumor nodule from the KP LucOS GEMM (expressing immunogenic neoantigens) showing H&E staining, a multiplexed CyCIF image of immune and tumor markers (DNA stain blue), RNAScope™ *in situ* hybridization for Cxcl9 and Cxcl10 (DNA stain blue), a map showing distance of each cell from the tumor edge (‘tumor boundary’), a map of cell type annotations, and a ‘graph’ map of physically interacting cells generated by Delaunay Triangulation showing the spatial organization of interacting cellular networks in and around tumor nodules. H&E, CyCIF and RNAScope™ images were acquired from serial sections. Maps of cells were derived from the CyCIF data. (C) Single cell gallery of lineage, cell state, and functional markers from representative, CyCIF images of the KP LucOS GEMM. (D) Sequential clustering of processed CyCIF imaging data using the marker combinations outlined in **Figure S1C** to define immune, epithelial tumor, and stromal cell populations. Rows represent individual cells.

The resulting data were analyzed using several computational approaches. For CyCIF, images were stitched and registered and then segmented to identify single cells (typically ∼100,000 to 500,000 cells per sample/mouse) and staining intensities quantified at the single-cell level; for mRNA ISH, foci were identified, and their density quantified, and the data were registered to CyCIF data from serial sections (see Methods). Distance metrics were used to characterize the position of cells relative to the boundary between tumor nodules and non-neoplastic lung tissue (i.e., the tumor edge) and to blood vessels (**Figure 1B**). The positions of single cells were then used to identify interacting cells in physical proximity and to create ‘graphs’ of interacting cell “networks’’ (**Figure 1B**).

### Tumor neoantigen expression reorganizes the immune landscape in KP lung cancer

We first profiled immune responses triggered by the LucOS CD8 T-cell neoantigens at a time point characterized by a transition between a functional and dysfunctional CD8 T-cell response (Burger et al., 2021; DuPage et al., 2011; Schenkel et al., 2021). At this time point (8 weeks after lentiviral infection), the tumor burden in KP LucOS mice is significantly lower than KP Cre mice (**Figure 2A**). However, the introduction of immunogenic CD8 T-cell neoantigens resulted in only modest differences in immune cell composition when lung tissue was examined as a whole (i.e., both tumor and non-tumor compartments together). For example, the numbers of neutrophils and B cells were slightly higher in KP LucOS whole lungs as compared to KP Cre lungs whereas Treg cells and dendritic cells were slightly lower, but these differences did not reach statistical significance (**Figures 2B-2D**).

**Figure 2.**
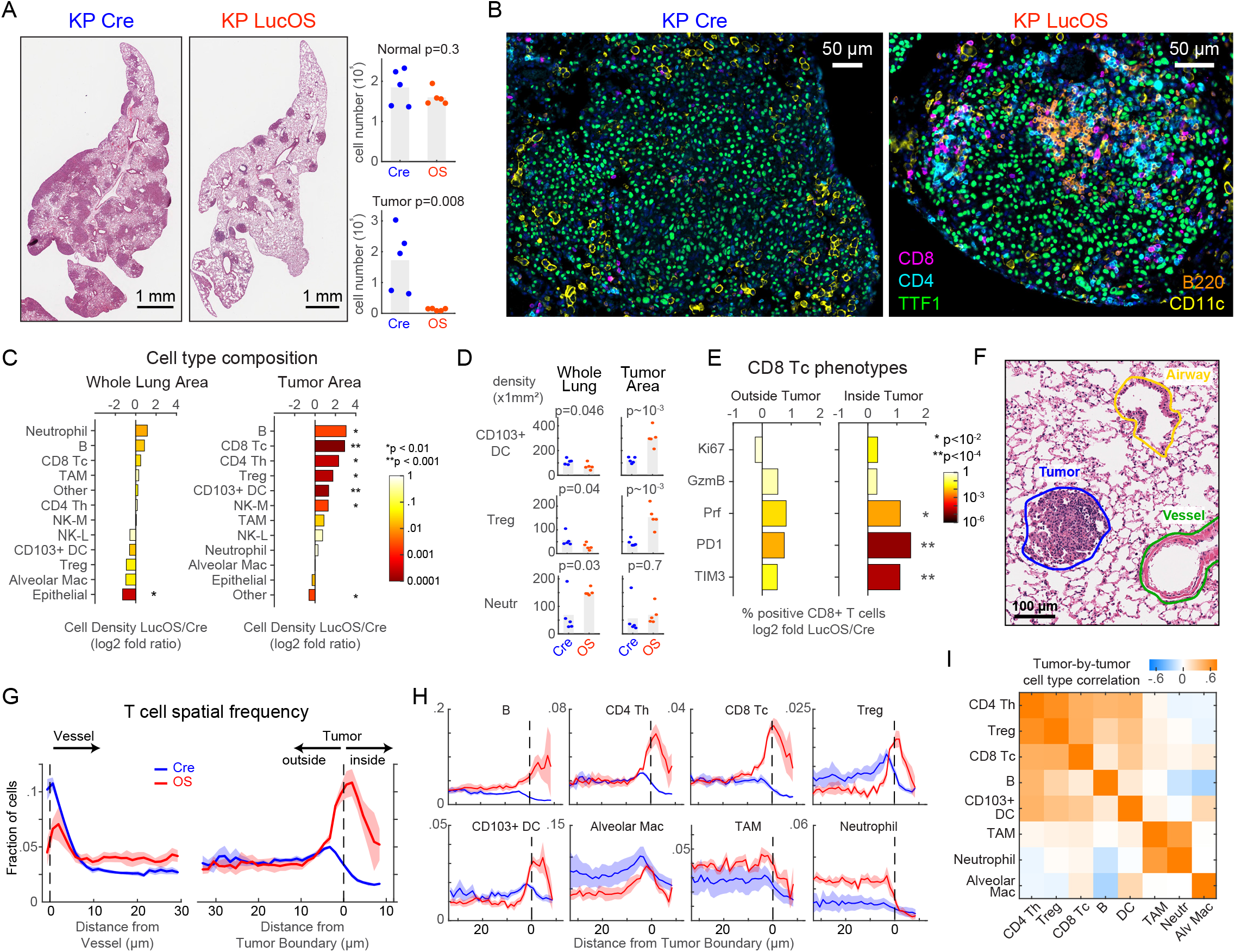
Tumor neoantigen expression reorganizes the immune landscape in KP lung cancer. (A-B) Representative H&E (A) and CyCIF (B) images of KP Cre versus KP LucOS (neoantigen-expressing) tumors and quantification of normal and tumor cell number (n = 5 mice per group, bar = mean). (C) Log2 fold ratio of cell-type densities between LucOS and Cre in whole lung areas and tumor areas (n = 5 mice per group, color represents two tailed t-test p-value). (D) Cell density measurements for indicated immune cell types in whole lung areas and tumor areas (n = 5 mice per group, bar = mean). (E) Log2 fold ratio between LucOS and Cre of density CD8 T-cell positive for single phenotypic markers indicated (right, inside tumor areas; left, outside tumor areas, n = 5 mice per group). (F) Representative image of pathology annotation of H&E showing tumor, airway, and blood vessel. (G) Probability density plots of T-cell spatial frequency relative to blood vessels and tumor boundaries in Cre and LucOS mice (n = 5 mice per group, mean +/- SEM). (H) Probability density plots of the frequency of indicated cell types from tumor boundaries in Cre and LucOS mice (n = 5 mice per group, mean +/- SEM). (I) Plot of tumor-by-tumor spatial correlation values within LucOS tumor nodules for indicated immune cell types (n = 29 tumors).

By contrast, when tumor areas were examined separately from non-neoplastic areas, the density of all lymphocyte subsets (Tc, Th, Treg, and B cells) was significantly higher in LucOS tumors as compared to Cre tumors, an increase of 3.3 to 8-fold (**Figures 2B-2D**). Increased infiltration of LucOS tumors was observed even for Treg cells that were less abundant in KP LucOS as compared to KP Cre lung as a whole (>3-fold higher in LucOS versus Cre tumors) (**Figure 2D**). Both NK (myeloid lineage marker-defined, see **Figure S1C**) and dendritic cells were also significantly increased within LucOS tumors but not in whole lung tissues (**Figures 2C, 2D** and **S2A**). Notably, the ratio of Tc cells to Treg cells was significantly increased in LucOS tumors (5.8-fold; and to a lesser extent in non-tumor tissue) (**Figure S2B**), a hallmark of a more immune permissive tumor microenvironment (TME) (Facciabene et al., 2012). Additionally, Tc cells inside tumors were enriched for expression of the cytotoxicity-associated marker Prf and the inhibitory receptors PD1 and TIM3, suggestive of a greater functional anti-tumor response moving toward exhaustion (**Figure 2E**). Interestingly, FACS analysis of T-cell populations from dissociated tumor-bearing lung lobes from the same mice was consistent with the whole lung area analysis; no significant changes in Tc, Th or Treg populations were observed, but trends toward increased Tc cells and decreased Treg cells resulted in an increased Tc/Treg ratio (**Figure S2C**). Hence, we find that whole lung area analysis of T cells by dissociative techniques does not fully capture tumor-specific phenotypes.

To determine the effect of neoantigen expression on the spatial distribution of immune cells relative to blood vessels and the tumor margin, we combined CyCIF with anatomical annotations from H&E images (**Figures 2F** and **S1A**). In both Cre and LucOS samples, we observed lymphocytes accumulated near blood vessels, a pattern of perivascular accumulation that is also observed in human tumors (**Figure 2G**) (Pullamsetti et al., 2017). Lymphocytes in KP Cre animals were excluded from tumors, whereas in KP LucOS animals, the lymphocytes breached the tumor boundary and infiltrated into the tumor (**Figures 2B, 2G** and **2H**). Moreover, the degree of infiltration by different types of lymphocytes (i.e., B, CD4 Th, CD8 Tc, Treg cells) was highly positively correlated in individual tumor nodules (**Figure 2I**), suggesting coordinated infiltration into tumors. In contrast, most types of myeloid cells were evenly distributed in the normal lung tissue, without evidence of perivascular accumulation. Myeloid cells were more abundant at the tumor margin but did not infiltrate into tumors in either KP Cre or KP LucOS mice with the exception of dendritic cells, which readily infiltrated the tumor in the KP LucOS model with spatial patterns similar to lymphocytes (**Figures 2D, 2H** and **2I**). Tumor exclusion was particularly evident in the case of neutrophils, which were substantially more abundant in KP LucOS than KP Cre lungs (**Figure 2H**).

### Neoantigen expression is associated with intratumoral localization of lymphocyte networks (‘lymphonets’)

The co-occurrence of different types of lymphocytes in KP LucOS tumors (**Figure 2I**) prompted us to look for evidence of lymphocyte cell-cell interactions. We applied the Visinity method recently developed by our group (Warchol et al., 2022) to interactively identify and quantify spatial arrangements among cells in whole-slide tissue images (see Methods). This method organizes cells into a 2-dimensional (2D) embedding based on the cell types within a neighborhood of defined diameter (50 µm in this analysis); cells close to each other in this representation are surrounded by similar cell types (**Figure 3A**). When applied to the nearly ∼2.6 million cells in the combined datasets from Cre and LucOS mouse lungs, the shared embedding space revealed a clear separation of neighborhood composition associated with normal lung and tumor (**Figures S3A-S3C**). Notably, the lymphoid population accumulated in two areas of the plot (clusters) at the intersection of normal and tumor neighborhoods and encompassed both B and T cells (**Figures 3B** and **S3A-S3C**), quantitatively demonstrating the spatial coordination of lymphocytes within cellular neighborhoods.

**Figure 3.**
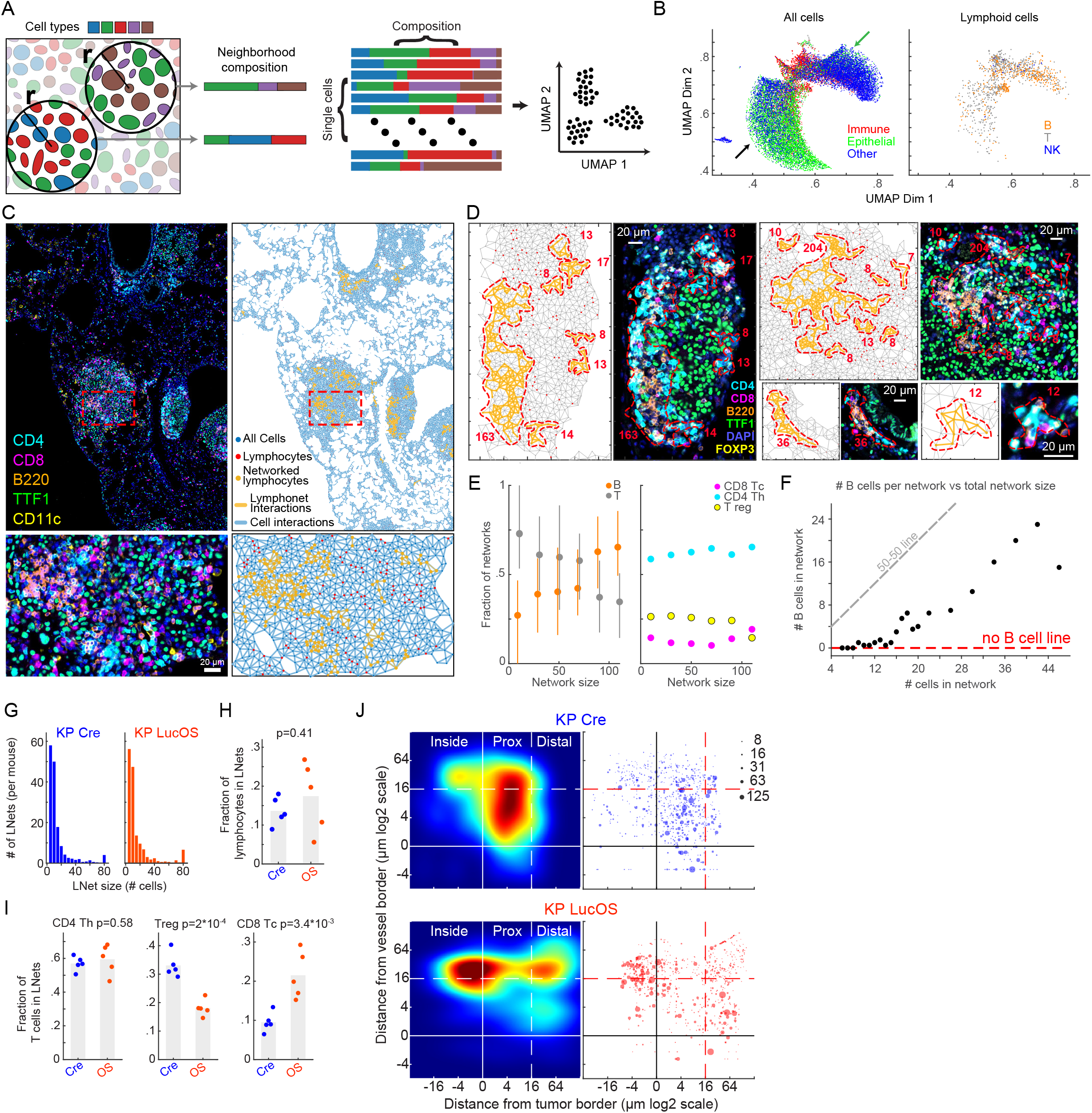
Neoantigen expression is associated with intratumoral localization of lymphocyte networks (‘lymphonets’) (A) Schematic diagram of neighborhood quantification using Visinity. Each cell within a given sample is assigned to a unique neighborhood. The neighborhood is defined as all cells within a specified radius to the reference cell. A feature vector is calculated based on this neighborhood which represents the weighted presence of each cell type within the given neighborhood. Groups of similar neighborhood vectors correspond to spatial patterns. (B) Neighborhood embedding generated by Visinity analysis of KP Cre and KP LucOS lung tissue with distinct regions highlighted that correspond to immune neighborhoods enriched in normal lung regions (green arrow) and others enriched in areas of tumor (black arrow). (C) CyCIF images and corresponding graphic maps of interacting immune populations identified by Delaunay Triangulation in KP LucOS lung tissue. (D) Examples of lymphonets of different sizes. (E) Lymphonet composition by cell type across various network sizes. Left proportion of B and T cells. Right, proportion of T-cell subtypes within T cell compartment (mean +/- 25^th^ percentile). (F) Scatter plot of number of B cells per network versus the total size of lymphonets. (G) Histogram of the number of lymphonets identified per mouse of indicated size in Cre and LucOS lung tissue. (H) Fraction of B and T lymphocytes and (I) T-cell subsets in lymphonets in Cre versus LucOS (n = 5 mice per group, bar = mean). (J) Left, density plots of lymphonets by distance from closest blood vessel (y-axis) and tumor (x-axis) in KP Cre and KP LucOS cohorts. Right, scatter plot lymphonets used to generate density plot (dot size represents the lymphonet size; n = 5 mice per group).

To characterize these T- and B-cell clusters, we generated graphs of cell-cell interactions by performing Delaunay Triangulation (Delaunay, 1934; Liebling and Pournin, 2012) (**Figures 3C-3D**; see Methods for computational details) on each specimen individually. We identified lymphocyte cell-cell networks that ranged from small clusters of less than ten lymphocytes to well over one-hundred lymphocytes that were in direct contact (**Figures 3C, 3D**; **Figure 3D** depicts examples of lymphonets ranging in size from 8 to 204 cells). Across KP Cre and LucOS mice, a minority of lymphocytes were organized into lymphonets (defined as ≥ 6 lymphocytes connected by direct cell-cell contacts; mean 15.5% ± 6.8% standard deviation (**Figure S3D**). We detected an average of ∼77 lymphonets per mouse lung lobe with an average of 17 cells per network. Analysis of lymphonet composition indicated that Th and B cells were the core structural components of lymphonets; over 60% of individual lymphonets had a majority of either Th or B cells (34% and 27%, respectively) in contrast to 5% having a majority of Tc cells and 8% containing a majority of Treg cells (**Figure S3E**). The fraction of B and T cells was strongly correlated with lymphonet size, with small lymphonets being enriched in T cells and large lymphonets being enriched in B cells (**Figure 3E**). Notably, lymphonets having fewer than 16 cells were almost exclusively composed of T cells and the frequency of B cells increased linearly after this threshold (**Figure 3F**). This relationship between network size and cell composition suggests that lymphonets nucleate from a core of T cells and subsequently grow by recruiting B cells.

While the overall number and size of lymphonets did not change substantially with neoantigen expression (KP Cre vs KP LucOS) across the lung tissues (**Figures 3G** and **3H**), lymphonets in KP LucOS contained significantly more Tc cells and significantly fewer Treg cells compared to lymphonets in KP Cre lungs (**Figure 3I**). In addition, neoantigen expression dramatically relocalized lymphonets relative to histopathological features (**Figure 3J**): in KP LucOS lungs, the majority of lymphonets were located inside tumors whereas in KP Cre mice most lymphonets were located outside of tumors, with a substantial fraction residing within 20 µm of a major blood vessel (**Figure 3J**). These findings reveal a strong correlation between neoantigen expression and lymphonet formation inside tumors.

### CXCR3 ligands modulate lymphonet formation and size but not intratumoral localization

The recruitment of activated Th and Tc cells to the TME is mediated in part by binding the CXCL9 and CXCL10 chemokines (and also CXCL11 in human) to the CXCR3 receptors on T cells (Metzemaekers et al., 2017; Mikucki et al., 2015). Given that small lymphonets predominantly contained T cells (**Figures 3E, 3F**), we hypothesized that CXCR3-mediated recruitment of T cells may contribute to the nucleation of lymphonets. Because CXCL9 and CXCL10 levels are tightly controlled at a transcriptional level (Ellis et al., 2010), and antibodies suitable for imaging cytokines in tissue are not available, we measured their distribution using RNA ISH (**Figures 1A, 1B**, and **4A**). In total, the levels of *Cxcl9* and *Cxcl10* mRNA in lung tissue were modestly increased in LucOS compared with Cre mice, but the changes were not statistically significant (**Figures 4B** and **S4A**). However, in KP LucOS (but not KP Cre) mice, *Cxcl9* mRNA expression was strongly localized within tumors, and both T and B cell localization was strongly spatially correlated with *Cxcl9* and *Cxcl10* expression (**Figures 4C**, **S4A** and **S4B**). In contrast, chemokine expression did not correlate with non-lymphocyte immune cells, such as neutrophils or tumor-associated macrophages (**Figure 4C**). Notably, the T and B cells that were spatially correlated with chemokine expression in LucOS mice were present within lymphonets (**Figures 4D** and **4E**).

**Figure 4.**
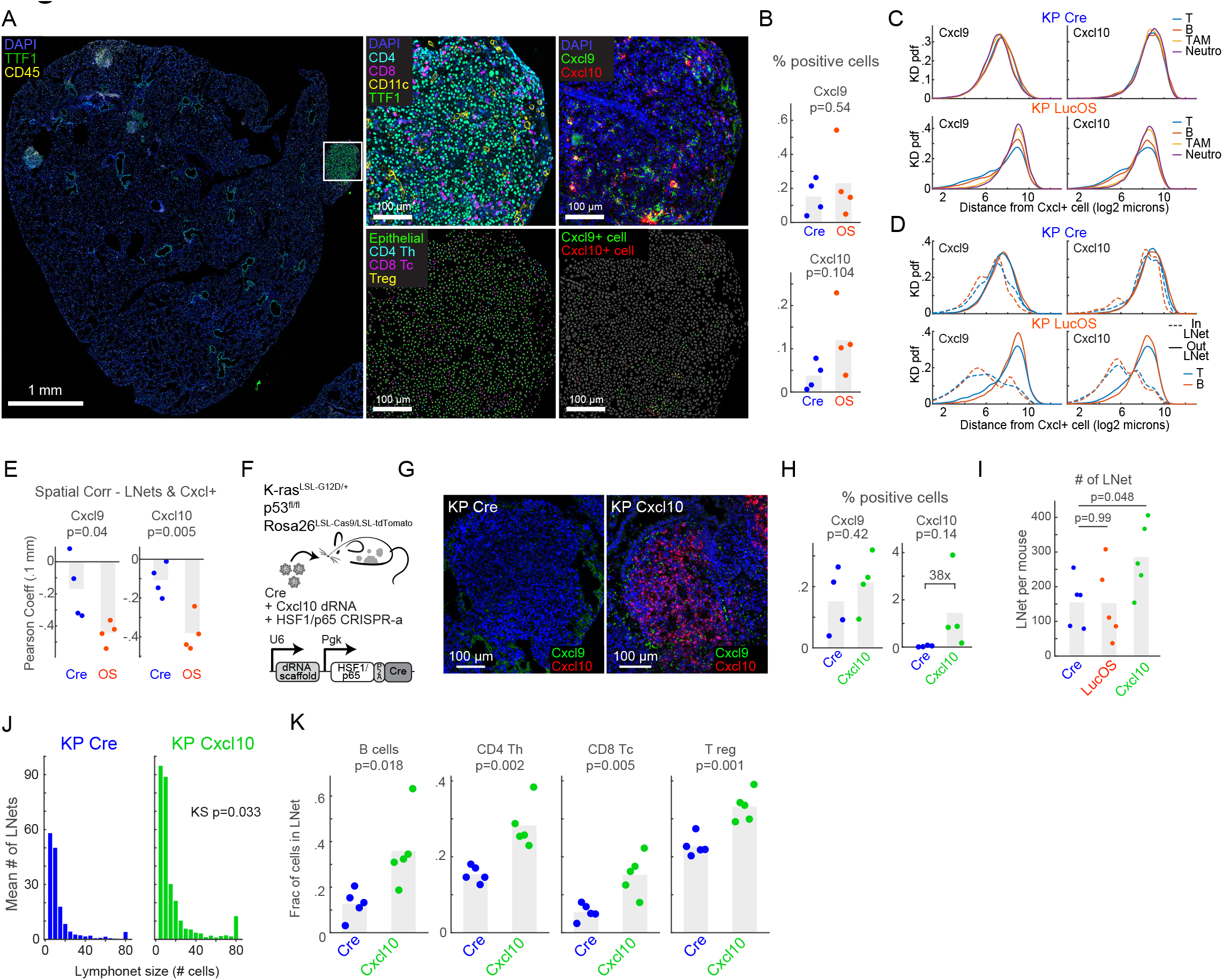
CXCR3 ligands modulate lymphonet formation and size but not intratumoral localization. (A) CyCIF and RNAScope™ in situ hybridization images from serial sections of a representative KP LucOS tumor nodule with cell type/state calls as indicated. (B) Percent of total cells expressing *Cxcl9* and *Cxcl10* mRNA in Cre versus LucOS lung tissue (n = 4 mice per group, bar = mean). (C-D) Kernel density probability density functions of the distance of the indicated immune cell populations (C) or T and B cells (D) in or out of lymphonets from *Cxcl9* and *Cxcl10* mRNA-expressing cells in Cre and LucOS. (E) Spatial correlation of *Cxcl9* or *Cxcl10* mRNA-expressing cells and lymphonets in Cre and LucOS (n = 4 mice per group, bar = mean). (F) Schematic of lentiviral system to deliver dRNAs and HSF1/p65 activation complex for CRISPR-a of *Cxcl10* in KP Cas9 mice. (G) Representative images of RNA *in situ* hybridization for *Cxcl9* and *Cxcl10* mRNAs using RNAscope™ in KP Cre versus KP *Cxcl10*-activated tumor nodules. (H) Percent of total cells expressing *Cxcl9* or *Cxcl10* mRNA in KP Cre versus KP *Cxcl10* lung tissue (n = 4 mice per group, bar = mean). (I) Number of lymphonets per mouse in KP Cre, KP LucOS, and KP *Cxcl10* (n = 5 mice per group, bar = mean). (J) Histogram of the mean number of lymphonets per mouse of indicated size in KP Cre and KP *Cxcl10* (n = 5 mice per group). (K) Plots of the fraction of indicated lymphocyte populations within lymphonets in KP Cre and KP *Cxcl10* (n = 5 mice per group, bar = mean).

To test whether CXCR3 ligands promote lymphonet formation, we used CRISPR-activation to ectopically express *Cxcl10* in KP Cre tumors (**Figure 4F**, see Methods); 38-fold induction of *Cxcl10* mRNA levels was achieved (**Figures 4G** and **4H**). Lymphonet number and size increased significantly in the lung (**Figures 4I** and **4J**) and involved recruitment of B cells and all T cell subsets (**Figure 4K**). Lymphonets were more proximal to blood vessels in KP mice over-expressing *Cxcl10* as compared to control KP Cre mice but remained markedly excluded from the inside of tumors (**Figure S4C**). These data show that expression of *Cxcl10* in the TME can promote the formation and growth of lymphonets but that additional neoantigen-dependent mechanisms are required for lymphonet localization to tumors.

### Spatial analysis reveals dynamic shifts in Tc cell states with immunotherapy treatments

To investigate the role played by lymphonets in anti-tumor Tc responses, we first assayed Tc function based on their expression of cytotoxic effectors (GzmB, Prf), checkpoint receptors (PD1 and TIM3), the TCF1 transcription factor, and the proliferation marker Ki-67. KP LucOS mice were exposed to one of two immunotherapy regimens previously shown to improve the anti-tumor functionality of the Tc response (Burger et al., 2021): (i) therapeutic vaccination (Vax) against the SIIN and SIY neoantigens, and (ii) antibody-mediated PD1/CTLA-4 immune checkpoint blockade (ICB) (**Figure S5A**). For Vax, KP LucOS mice were injected subcutaneously with SIIN and SIY 30-mer peptides and cyclic-di-GMP as an adjuvant at 6 weeks post-tumor initiation followed by a booster at 8 weeks and then sacrificed at 9 weeks. For ICB therapy, a mixture of anti-PD1 and anti-CTLA-4 antibodies or, as a control, isotypes controls were administered by injection into the peritoneal cavity starting at 8 weeks post-tumor initiation (a total of three doses spaced three days apart: day 0, 3 and 6 of week 8) and then the mice were sacrificed, also at 9 weeks after tumor initiation.

The resulting data were analyzed using Palantir, an algorithm that uses multidimensional expression data to align single cells along differentiation trajectories, thereby capturing continuity in cell states and stochasticity in cell fate determination (Setty et al., 2019). Three predominant CD8 T-cell states (S1 to S3, **Figures 5A** and **S5B**) were identified in both Vax and ICB mice and gated using a supervised method typical of FACS data analysis (see Methods). S1 had high levels of TCF1 expression and no expression of markers of activation/exhaustion (i.e., PD1, TIM3) or cytotoxicity (i.e., GzmB, Prf) and thus corresponded to a naïve T cell state (**Figures 5B** and **S5C**). S2 had high expression of GzmB and/or Prf and the proliferation marker Ki-67, indicative of a proliferative, cytotoxic T cell state. S3 had low expression of GzmB, Prf, and Ki-67 and high expression of the inhibitory receptors PD1 and TIM3, denoting an exhausted T cell state. These three states were interconnected by cells – about one-third of the total – having transitional phenotypes (T1, T2, T3) in which the expression of multiple markers was graded and mixed (**Figures 5A, 5B**, and **S5B-S5C**). Using this division of cell types and states, we examined shifts in CD8 T-cell function induced by the two immunotherapy regimens.

**Figure 5.**
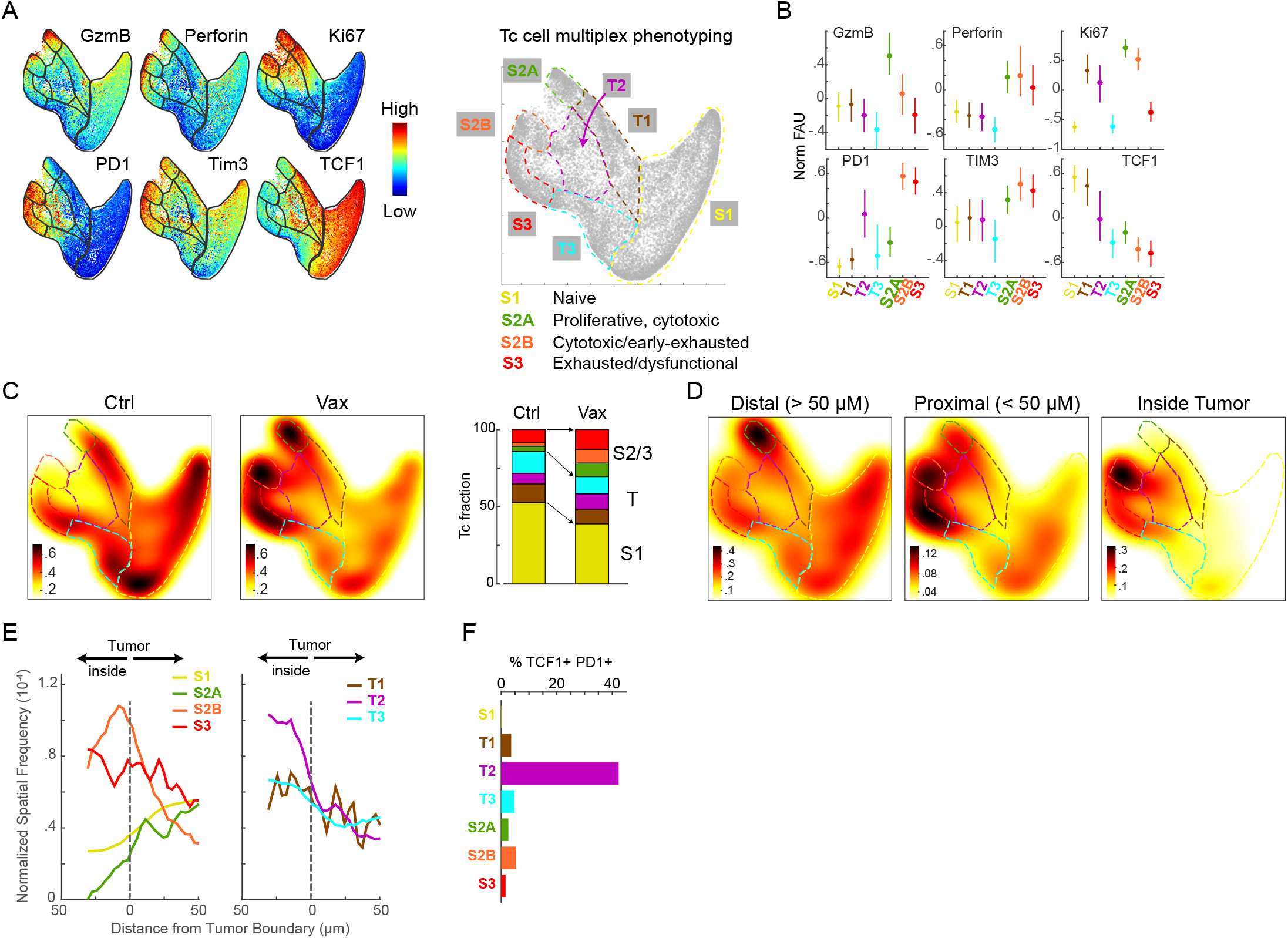
Spatial analysis reveals dynamic shifts in Tc cell states and localization with immunotherapy treatments. (A) Palantir projection of Tc populations in KP LucOS mice treated with a SIINFEKL (SIIN) and SIYRYYGL (SIY) long-peptide vaccine (Vax) or PBS (Ctrl) (n = 10^4^ cells sampled from n = 8 and 7 mice per treatment, see **Figure S5A** for treatment schematic). The expression levels of the indicated markers are mapped to color (normalized between 0.1 and 99^th^ percentile). Tc states defined by multiparameter measurements are indicated at the extremes of the representation (S1, S2A, S2B, and S3) connected by transitional phenotypes (T1-T3) shown in the schematic to the right. (B) Normalized fluorescence units for each of the markers in the indicated Tc cell states and transitions (mean +/- 25^th^ percentile). (C) Heat map of Tc cell densities in Palantir projections for Ctrl and Vax groups (n = 10^4^ cells per treatment). Right, stacked bar graph of the fraction of Tc cells in each state and transition. (D) Heat map of Tc densities in Palantir projections for KP LucOS mice following Vax separated by distance from tumor boundary (distal: >50 µm from the tumor boundary; proximal: <50 µm from the tumor boundary; and inside tumor). (E) Spatial frequency of Tc cell states and transitions from tumor boundaries (Vax). (F) Percent of Tc cells that are TCF1+ PD1+ in each Tc cell state.

In the untreated LucOS cohorts, the majority of Tc cells were naïve (S1), but Vax and ICB protocols shifted cells into cytotoxic (S2) and exhausted (S3) states (**Figures 5C** and **S5D**). In the Vax cohort, the cytotoxic (S2) population split into two groups distinguished by the levels of PD1 and TIM3 expression (S2A and S2B in **Figures 5A-5C**): the S2A state had low PD1/TIM3 expression and appeared highly cytotoxic, expressing high levels of both GzmB and Prf whereas cells in the S2B state expressed high levels of PD1/TIM3 cells and expressed GzmB at relatively lower levels. In the phenotypic landscape, cells in the S2B state were adjacent to the exhausted (S3) population, suggesting S2B may represent a cell state on the verge of exhaustion/dysfunction. In the ICB cohort, the S2 state did not split and resembled the PD1/TIM3^high^ GzmB^low^ state of S2B Vax cells (**Figures S5B-S5D**). These data suggest that Vax is substantially more effective than ICB in generating a highly cytotoxic and proliferative effector state lacking markers of exhaustion.

### Functionally distinct Tc cell states are spatially segregated in the tumor microenvironment

To characterize the spatial distribution of the CD8 T-cell states relative to tumor cells, we split the Palantir phenotypic landscape depending on whether the immune cells (i) resided inside tumors, (ii) were proximal (< 50 µm) to the edge of tumors, or (iii) were distal to tumors (>50 µm away from tumor edges) (**Figures 5D-5E** and **S5E-S5G**). Strikingly, we found that the highly cytotoxic S2A state that was unique to Vax mice was present distal to tumors (**Figures 5D** and **5E**), whereas the cytotoxic/early-exhausted S2B (Vax) and S2 (ICB) states were enriched inside tumors (**Figures 5D, 5E**, **S5E-S5F**). The exhausted S3 population in both Vax and ICB mice was found proximal to tumor edges, indicating that this dysfunctional state is associated with exclusion from tumors (**Figures 5D, 5E**, and **S5F-S5G**). These findings show that the spatial segregation of different functional states of CD8 T cells is altered in an immunotherapy-specific manner both within tumors and surrounding normal tissue.

Neither the Vax nor ICB protocols significantly changed the fraction (∼30%) of CD8 T cells that displayed transitional phenotypes (T1-T3; **Figures 5C** and **S5D**). In Vax, T2 cells were both spatially enriched inside tumors (**Figure 5E**) and were dramatically enriched for cells co-expressing TCF1 and PD1 (between 8- and 200-fold increase above other states, **Figure 5F**), similar to T1 and T2 states in ICB (**Figure S5H**). TCF1+ PD1+ CD8 T cells have recently been shown to play a critical role in driving therapeutic responses to ICB therapy in both mice and humans (Philip and Schietinger, 2022). Such cells are in a progenitor-like state and are induced to differentiate into cells having cytotoxic function in response to treatment (Kurtulus et al., 2019; Siddiqui et al., 2019). The findings from our combined spatial and phenotypic analysis suggest that TCF1+ PD1+ progenitor CD8 T cells are enriched in specific transitional states that efficiently traffic into tumors and can establish residence within the tumor bed.

### TCF1+ PD1+ progenitor CD8 T cells reside within intratumoral lymphonets

We next used the data from the Vax-treated LucOS mice to investigate how lymphonets intersect with the phenotypic landscape of Tc cells. While Vax treatment did not substantially change the overall size, number, or localization of lymphonets (**Figures S6A-S6B**), Vax increased lymphonet association of CD8 T cells but not other T cell subsets (**Figures 6A-6B**, and **S6C)**. Remarkably, the TCF1+ PD1+ progenitor CD8 T cell state was the most highly and significantly enriched state in lymphonets (**Figure 6C**, KS p-value = 10^-3^). Across the Vax cohort the total number of Tc cells in lymphonets was linearly correlated with the number of TCF1+ PD1+ cells (**Figure 6D**). Consistent with this, the CD8 T-cell compartment of lymphonets was predominantly comprised of the transitional T2 state containing TCF1+ PD1+ progenitor cells (**Figures 6E-6F** and **S6D**); this was true of lymphonets both inside and outside of tumors, however, cells in the T2 state were mostly found within tumors (**Figures 5D** and **5E**). Lymphonets were similarly enriched for the transitional phenotypes containing TCF1+ PD1+ cells in the ICB cohort (i.e., T1 and T2, **Figures 6G-6H** and **S6E**). Notably, the only CD8 T-cell state that increased in lymphonets following Vax or ICB treatment was the cytotoxic S2 state (S2B for Vax and S2 for ICB, **Figures 6E-6H**). Thus, after treatment, cytotoxic cells colocalized with TCF1+ PD1+ progenitor CD8 T cells in lymphonets. Given that TCF1+ PD1+ progenitor cells give rise to cytotoxic Tc cells in tumors (Siddiqui et al., 2019), these data suggest that lymphonets are the site of differentiation of progenitor cells into cytotoxic cells in response to immunotherapy.

**Figure 6.**
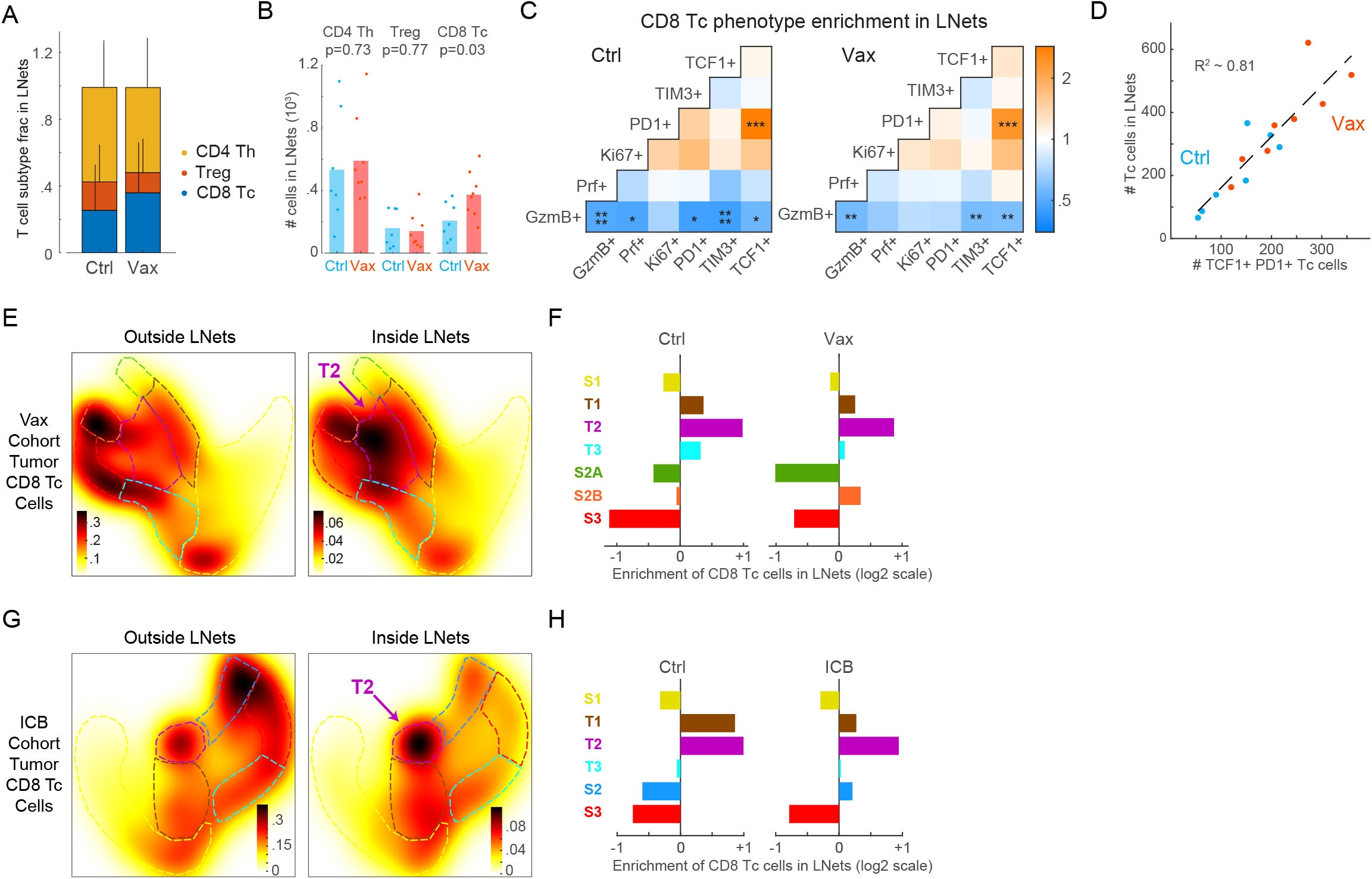
TCF1+ PD1+ progenitor CD8 T cells reside within intratumoral lymphonets. (A) Proportion of T-cell subtypes in lymphonets (Ctrl n = 7, Vax n = 8 mice, mean + SD, same KP LucOS cohort as Figure 5). (B) Number of cell T-cell subtypes present in lymphonets (bar = mean of n = 7 and 8 mice). (C) Pairwise enrichment analysis of marker co-expression in Tc cells in Ctrl and Vax groups (KS p-value *p =0.05, **p = 0.01,***p = 10^3^,****p = 10^4^. (D) Scatter plot of Tc cells present in lymphonets versus TCF1+ PD1+ cells in both Ctrl and Vax groups (dotted line, linear regression, R^2^ = 0.81). (E) Heat map of cell densities of tumor-localized Tc cells present outside and inside lymphonets in Palantir projections for Vax-treated cohort (n = 3,736 and 806 cells, respectively). (F) Enrichment of tumor-localized Tc cells in lymphonets for Ctrl (n = 7) and Vax (n = 8) mice. (G) Heat map of cell densities of tumor-localized Tc cells present outside and inside lymphonets in Palantir projections for anti-PD1 and anti-CTLA4 treated (ICB) cohort (n = 6 mice per group, n = 4,276 and 1,041 cells respectively). (H) Enrichment of tumor-localized Tc cells in lymphonets for Ctrl and ICB mice (n = 6 mice per group).

### Lymphonets enriched for TCF1+ PD1+ progenitor CD8 T cells are abundant in early-stage human lung adenocarcinoma

We used a panel of CyCIF-qualified antibodies to characterize the features of lymphonets in 14 whole slide sections of early-stage human lung adenocarcinoma (**Table S2**), to parallel the early-stage tumors studied in the KP LucOS GEMM. We performed sequential clustering of ∼7.8 million cells from these images to identify tumor and stromal cells (**Figure 7A**, Lv1) and to computationally isolate immune cells (∼3.4 million cells) for further cell-type calling (**Figure 7A**, Lv2-Lv4). The samples had highly variable fractions of each of the cell types and lymphocyte subtypes (**Figure 7B**). In histopathologically-annotated tumor areas, we identified many lymphonets per sample, and they varied substantially in size. Similar to lymphonets in mice, the vast majority of these networks in human tumors were small (**Figures 7C** and **7D**), and the fraction of B cells was positively correlated to lymphonet size (**Figure 7E**). Notably, only a few of these lymphocyte networks were similar to the organized aggregates of T and B cells that are known as tertiary lymphoid structures (TLS, **Table S3**) (Schumacher and Thommen, 2022), as scored by pathology review (see Methods). These findings suggest that the anti-cancer immune response in both human and early-stage mouse lung cancer is characterized by a preponderance of smaller lymphocyte networks. Similar to the KP mouse tumors, in the human lung tumors, T cells were prevalent in smaller lymphonets. Uniquely to human samples, the fraction of CD8 T cells decreased as lymphonets increased in size, being replaced by CD4 Th cells (**Figure 7E**). A positive spatial correlation between tumor cell major histocompatibility class I (MHC I) expression and lymphonets was observed (**Figure 7F**), suggesting lymphonet organization in early-stage human lung cancer may be regulated by CD8 T-cell antigen presentation. A negative spatial correlation was observed between tumor cells expressing PD-L1 and lymphonets (**Figure 7F**), suggesting that PD-L1 may support the exclusion of lymphonets. Similar to **Figure 5**, we profiled Tc cells with functional markers and used Palantir to identify the TCF1 and PD1 co-expressing population of progenitor CD8 T cells. While Tc cells were present both outside and inside of lymphonets, TCF1+ PD1+ progenitor cells were largely restricted to lymphonets (**Figure 7G**) and became increasingly enriched as lymphonet size increased (**Figure 7H**). Altogether, these findings reveal that lymphonets as defined in the KP GEMM model are found in abundance in human lung adenocarcinomas and may similarly support progenitor CD8 T-cell maturation.

**Figure 7.**
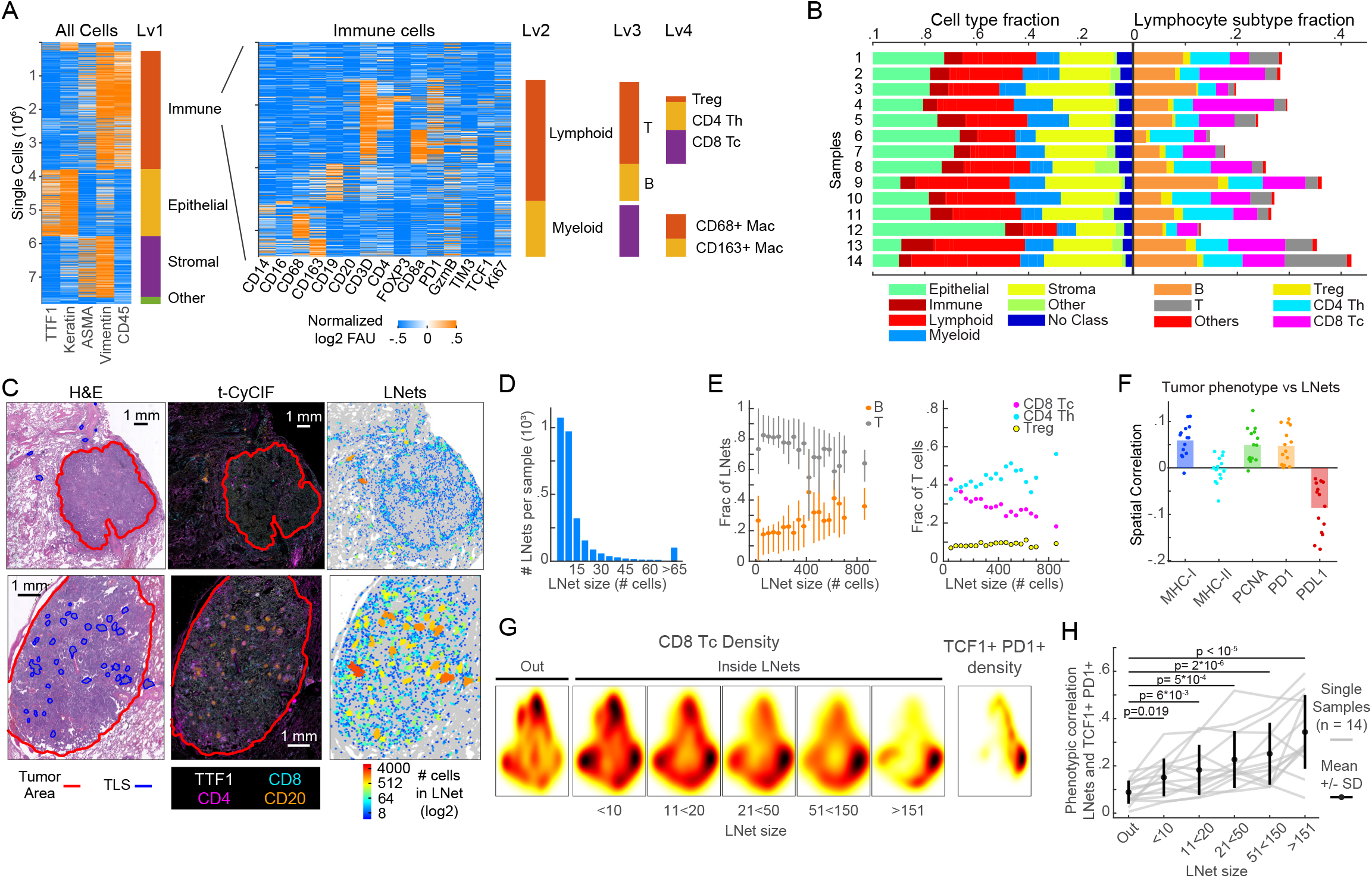
Lymphonets enriched for TCF1+ PD1+ progenitor CD8 T cells are abundant in early-stage human lung adenocarcinoma. (A) Sequential clustering of immune, epithelial tumor, stromal and ‘other’ cell populations (Lv1); the immune cells were further clustered into lymphoid and myeloid cells (Lv2) and immune subsets (Lv3 and Lv4: Treg, CD4 Th, CD8 Tc, B, CD68+ macrophages, CD163+ macrophages). Rows represent individual cells. 7.8 x10^6^ cells are plotted from n = 14 human lung adenocarcinomas (**Table S2**). Immune clusters are shown in the heat map at the right. (B) Horizontal stacked bar graphs of cell type fractions (Lv1-2) and lymphocyte subtype fractions (Lv3-Lv4). (C) Representative images of H&E, CyCIF, and maps of lymphonets colored to indicate size in two examples of human adenocarcinoma. Top, exemplar case of tumor presenting overwhelmingly with small lymphonets (n<64 cells). Bottom, exemplar case of tumor presenting with large lymphonets (n>64 cells). (D) Histogram of the mean number of lymphonets per sample (n = 14) binned by lymphonet size (5 to >65 cells). (E) Composition of lymphonets by cell type (B vs. T; CD8 Tc vs. CD4 Th vs. CD4 Treg) across different network sizes (mean +/-25^th^ percentile). (F) Spatial cross-correlation of lymphonets and expression of indicated markers (n = 14 samples, bar = mean). (G) Heat map of density of total CD8 T cells in and out of lymphonets of different sizes, and density of TCF1+ PD1+ CD8 T cells in Palantir projection from 14 human lung adenocarcinomas (n = 10^4^ cells sampled from n = 14 samples). (H) Spatial correlation of TCF1+ PD1+ CD8 T cells to lymphonets binned by lymphonet size; individual gray lines represent data from single human lung cancers (n = 14) and black line is the mean ± standard deviation.

## DISCUSSION

Recent advances in single-cell and multiplexed spatial analysis of tissues are driving large scale inter-institutional and international efforts to systematically characterize the spatial organization of human tumor and immune cells during cancer development and in response to therapy (Rajewsky et al., 2020; Rozenblatt-Rosen et al., 2020). Recurrent spatial features of human cancers are being identified, for example i) active immune surveillance of proliferating tumor cells by cytotoxic T cells (Gaglia et al., 2022; Launonen et al., 2022), ii) chemokine-organized tumor-immune architecture (Nirmal et al., 2022; Pelka et al., 2021), and iii) and cell-cell interactions involved in generating extracellular metabolites such as adenosine, an immunosuppressive purine metabolite of ATP hydrolysis by the CD39 and CD73 ectoenzymes (Coy et al., 2022). These early insights into the tissue cellular neighborhoods and cellular architectures of human cancer tissues have revealed a pressing need for experimental and analytical approaches to investigate the mechanistic underpinnings of the anti-cancer immune response.

Cancer GEMMs are a natural complement to analysis of human tumor samples because they provide a highly physiological and tractable experimental system for deep mechanistic study of the regulators of the tumor-immune microenvironment (Connolly et al., 2022; Dhainaut et al., 2022; Wroblewska et al., 2018). However, substantial method development is required to perform high-dimensional spatial characterization of the tumor-immune interactions in these models. In this study, we describe an approach for spatial interrogation of the immune microenvironment of genetically engineered mouse models (GEMMs) of cancer by imaging mRNA and protein expression and integrating this information with pathology-annotations of tissue landmarks acquired from conventional H&E stains. This work shows that the composition, functional states, and spatial organization of the immune response to neoantigen expression and immune therapies can be efficiently mapped using multi-marker measurements from whole lung mouse tissues. The CyCIF panels and the computational analysis pipelines used here are available in the public domain and can be applied across distinct cancer histologies using conventional microscopes and widely available reagents (Lin et al., 2018; Schapiro et al., 2022a, 2022b). We expect that the antibodies and analysis methods developed in this effort can also be adapted for use with other multiplexed imaging methods that are in use to acquire multiplexed tissue images.

Spatial characterization of the KP GEMM of lung cancer revealed striking differences that are induced by tumor neoantigen expression. These changes in the immune landscape were observed in the tumor areas but were not identified when analyzing entire lung areas, underscoring the limitations of whole-tissue dissociative methods for faithfully capturing localized or tumor-specific phenotypes. Comparison of immunogenic (KP LucOS) versus non-immunogenic (KP Cre) tumor-bearing lungs revealed no significant differences in immune cell composition across whole lung areas but pronounced differences in lymphocyte and dendritic cell localization to tumors. KP LucOS tumors heavily recruited T and B cells in a coordinated manner and formed lymphoid networks ranging from as few as 6 cells to upwards of hundreds of cells that we termed lymphonets, predominantly composed of Th cells and B cells, and harboring ICB-responsive progenitor TCF1+ PD1+ CD8 T cells. The ability to introduce genetic and therapeutic perturbations in the KP GEMM allowed us to interrogate the regulation of lymphonet formation and the role of lymphonets and the spatial segregation of CD8 T cell subsets in coordinating anti-tumor immunity.

Multiparametric analysis of key functional Tc cell markers in KP LucOS tumors using the Palantir algorithm defined three major Tc cell states, naïve (S1), cytotoxic (S2), and dysfunctional/exhausted (S3), and characterized the flux through these states and connecting transitional phenotypes (T1-T3) in response to immunotherapies. Neoantigen vaccination (Vax) and anti-PD1/anti-CTLA-4 immune checkpoint blockade (ICB) shifted Tc cells from the naïve S1 state to the S2 and S3 states. These differentiated functional states were connected by cells exhibiting intermediate transitional phenotypes (T1-T3). TCF1+ PD1+ cells that have been described as giving rise to cytotoxic and exhausted CD8 T-cell populations in response to ICB therapy (Philip and Schietinger, 2022) occupied the intratumoral transition states and were tightly associated with lymphonets both before and after immunotherapy treatment. After therapy, S2 cells colocalized with TCF1+ PD1+ cells in lymphonets, consistent with progenitor cells seeding the S2 population. Notably, vaccination resulted in two S2 populations (highly cytotoxic S2A T cells marked by high expression of GzmB, and cytotoxic/early-exhausted S2B T cells marked by low expression of inhibitory receptors) that were spatially segregated; only the S2B population localized to tumors and lymphonets while the S2A population was present outside of tumors. The exhausted/dysfunctional T cells (S3) were excluded just outside of the tumor margin. We hypothesize that in contrast to the S2B (and ICB S2 populations), S2A cells are not derived from intratumoral TCF1+ PD1+ cells and instead seed directly from the periphery. Upon entering tumors, S2A cells may pass through the S2B state before they become terminally exhausted (S3). Consistent with this, we previously reported that vaccination acutely promotes substantial peripheral CD8 T-cell expansion rather than expanding the existing CD8 T-cell populations in the lung by flow cytometric analysis (Burger et al., 2021). In contrast to Vax, ICB induced only the intratumoral S2B-like S2 state associated with TCF1+ PD1+ progenitor cells, and this may help to explain the central role of progenitor cells in driving ICB response in mice and humans.

Compartmentalized and structured rather than mixed organization of lymphocytes with respect to tumors has previously been correlated with tumor control (Keren et al., 2018), particularly the formation of tertiary lymphoid structures (TLS) observed across multiple cancer types (Colbeck et al., 2017; Schumacher and Thommen, 2022). TLS are aggregates of immune cells with cellular composition and organization resembling secondary lymphoid organs, with fully mature TLS generally described as having B- and T-cell zones and germinal centers, containing follicular dendritic cells, and being vascularized by high endothelial venules (HEVs). The presence of TLS is predictive of better patient survival and response to immune checkpoint blockade and vaccine immunotherapies across multiple cancer types (Cabrita et al., 2020; Helmink et al., 2020). However, it remains unclear whether TLS directly facilitate anti-tumor immune responses or are merely evidence of a prior immune response with potential for reinvigoration by immunotherapy. Characterization of dynamic changes within TLS over time or with therapy is difficult to investigate in humans and studies in mice have been limited due to the absence of TLS formation in most transplantable tumor models (Fridman et al., 2022). In the KP LucOS model, we previously described the formation of mature TLS peritumorally around 20 weeks post-tumor initiation (Joshi et al., 2015), a time-point correlated with loss of a functional anti-tumor CD8 T-cell immunity and lack of response to anti-PD1/anti-CTLA-4 ICB therapy (Burger et al., 2021; DuPage et al., 2011; Schenkel et al., 2021). In comparison to these TLS, the lymphonets we observed (at 9 weeks post-tumor initiation) in conjunction with functional CD8 T-cell responses were localized inside tumors and were less structured, lacking distinct T- and B-cell zones and significant association with dendritic cells. It is possible that some lymphonets represent precursors to the TLS observed later during tumor progression. Further spatial profiling of the tumor microenvironment longitudinally between 9- and 20-weeks post-tumor initiation could shed light on this possibility and identify factors regulating a shift from lymphocyte structures supporting CD8 T-cell immunity to those having bystander or immunosuppressive function.

Consistent with our observation in mice that intratumoral lymphonets harbor TCF1+ PD1+ progenitor CD8 T cells, we found that TCF1+ PD1+ cells were also localized to lymphonets in human lung cancer resections. Similarly, localization of stem-like cells (defined as CXCR5+ TCF1+) to intratumoral lymphocyte ‘niches’ has been previously reported in human renal cell carcinoma, where the ‘niches’ were proposed to support generation of cytotoxic T cells (Jansen et al., 2019). These niches were not mature TLS and instead were defined by lymphocyte aggregation around MHC II-expressing cells, presumably marking regions rich in antigen presenting cells. Interestingly, we did not find a correlation between MHC II expression and lymphonets of any size in human lung cancer or a significant association of dendritic cells with lymphonets in mouse or human. However, MHC I expression level was correlated with lymphonets in human tumors; this may suggest that antigen presentation to CD8 T cells is necessary for lymphonet formation and/or that lymphonets promote MHC I upregulation (perhaps through T cell secretion of IFNɣ). Consistent with this observation, lymphonets in the mouse were found intratumorally only with expression of LucOS neoantigens and these lymphonets were significantly associated with cells expressing IFNɣ-induced chemokines (i.e., CXCL9 and CXCL10). Interestingly, we observed that ectopic expression of CXCL10 was able to increase the size and number of lymphonets in Cre mice lacking neoantigen expression. Pelka et al. recently reported a significant association between formation of “immune hubs” enriched in T lymphocytes (similar to the lymphonets reported here) and expression of CXCR3 ligands during a productive anti-tumor immune response to mismatch repair deficient (MMRd) human colorectal cancer (Pelka et al., 2021). Our findings provide mechanistic evidence suggesting CXCR3 ligands actively promote the formation of lymphocyte niches correlated with productive anti-tumor immunity; however, localization of these cell networks inside tumors depends on neoantigen expression or associated factors.

Lymphonets in KP LucOS mice were predominantly composed of Th and B cells, with the fraction of B cells increasing with lymphonet size in both mouse and human. Notably, an association of B-cell gene signatures with better patient survival and response to ICB therapy has been found with remarkable frequency across many cancer types (Fridman et al., 2022). Interestingly, however, B cells in cancer have been demonstrated to have both pro- and anti-tumorigenic functions. For example, B regulatory (Breg) cells contribute to tumor-promoting inflammation and suppression of anti-tumor T-cell responses, while antibody-producing plasma cells (frequently associated with TLS) are more commonly associated with tumor control (Fridman et al., 2022). Future spatial proteomics studies with additional markers of B-cell states paired with spatial transcriptomics in the KP GEMM could clarify the function of B cells and Th cells in lymphonets and how they might support TCF1+ PD1+ progenitor CD8 T-cell function. Given that antigen is necessary for nucleation of lymphonets inside KP lung tumors and MHC I expression is associated with lymphonets in human lung cancer, one hypothesis is that B cells regulate CD8 T cells and support Th-cell function through their role as antigen presenting cells (Bruno et al., 2017).

## Limitations of this study

One challenge with this study was that the antibody panel we qualified was largely focused on phenotyping effector T-cell states, but additional antibodies will be needed to adequately characterize other important T-cell populations (i.e., resident memory, T follicular helper cells) as well as the diversity of tumor-associated myeloid cells that can be monitored using scRNA-seq methods (Leader et al., 2021). Use of emerging spatial transcriptomics methods may soon facilitate deeper phenotyping of some immune cell states and expand the range of cytokines and chemokines that can be simultaneously profiled (Dhainaut et al., 2022). Another challenge was that the MHC-peptide tetramers that are commonly used to identify antigen-specific T cells in FACS analysis are not active in FFPE tissue sections, thus, we were not able to detect and quantify the CD8 T cells that are specific for SIIN and SIY CD8 T-cell antigens. Being that many lymphonets largely comprise Th cells, additional analysis will be needed to understand the role of diverse neoantigens on lymphonets as well as the effects of CD8 T-cell antigens with different MHC affinity (Burger et al., 2021). Depleting immune populations (Hiam-Galvez et al., 2021) will also be useful for dissecting the contributions of individual cell types on lymphonets formation and function. An additional limitation is that multiparametric measurements permit the inference of dynamic properties, but they do not allow the direct visualization of transitions over time in the way that intravital imaging methods permit (Jain et al., 2002; Mueller et al., 2021); however, such microscopy methods image relatively restricted regions of tissue over short windows of time in small numbers of animals and the methods are prone to motion artifacts when applied to lung tissues. A considerable limitation of the current work is the limited availability of software and computational methods for processing and analyzing multiplexed tissue images. While our study leverages recent tool development and presents an analysis framework for quantifying networks of lymphocytes, the analysis of whole-slide high-plex tissue images is still at an early stage. Releasing full resolution Level 3 images and spatial feature tables (Schapiro et al., 2022a, 2022b) derived from the high-plex images from our study should accelerate tool and method development and advance the use of these tools for mechanistic studies of mouse models of cancer.

## ACKNOWLEDGEMENTS

This work was supported by NIH grants R01-CA194005 (S.S.), R41-CA224503 (P.K.S.), U54-CA225088 (P.K.S., S.S.), T32-GM007748 (S.C.), and T32-HL007627 (G.G.), by the Bridge Project, a partnership between the Koch Institute for Integrative Cancer Research at MIT and the Dana-Farber/Harvard Cancer Center (P.K.S., S.S., T.J.), the Howard Hughes Medical Institute (T.J.), the Ludwig Center at Harvard (P.K.S., S.S.), the American-Italian Cancer Foundation postdoctoral fellowship (G.G.), K99-CA256497 (A.J.N), fellowship awards from the Jane Coffin Childs Memorial Fund for Medical Research and the Ludwig Center for Molecular Oncology at MIT (M.L.B.), and the BWH President’s Scholar Award (S.S.). We thank Dana-Farber/Harvard Cancer Center for the use of the Specialized Histopathology Core, which provided histopathology services. Dana-Farber/Harvard Cancer Center is supported in part by an NCI Cancer Center Support Grant P30-CA06516. This work was supported in part by the Koch Institute Support (core) Grant P30-CA014051 from the National Cancer Institute. T.J. is a Daniel K. Ludwig Scholar.

## AUTHOR CONTRIBUTIONS

Conceptualization: G.G., M.L.B., P.K.S., T.J., S.S.; Methodology: G.G., M.L.B., S.W., A.M.J., S.N., A.J.N., R.K.; Data acquisition: G.G., M.L.B., C.C.R., D.R., Y.D., G.E.C., S.Z.T, A.M.J., S.N., S.C., J.R.L.; Software: G.G., S.W., A.J.N., R.K., H.P.; Validation: G.G., M.L.B., C.C.R., D.R., Y.D.; Formal Analysis: G.G., M.L.B., C.C.R., D.R., Y.D.; Resources: H.P., P.K.S., T.J., S.S., Data Curation: G.G., M.L.B., C.C.R., D.R., Y.D.; Writing – Original Draft: G.G., M.L.B., S.S.; Writing – Reviewing & Editing: all authors; Supervision: H.P., P.K.S., T.J., S.S.; Project Administration:; G.G., M.L.B., R.K., H.P., P.K.S., T.J., S.S.

## STAR METHODS

### Human Tissue

Formalin fixed paraffin embedded (FFPE) tissue samples of human lung adenocarcinoma were retrieved from the archives of the Brigham and Women’s Hospital Department of Pathology following approval of the research study by the Partners Healthcare Institutional Review Board at Brigham Health, Boston, MA, USA (Excess tissue, discarded tissue protocol number 2018P001627). All appropriate ethical guidelines were followed for this study.

### Mice

Lung adenocarcinomas were initiated in *Kras*^LSL-G12D/+^; *p53*^fl/fl^ (KP) on a C57BL/6 background through intratracheal installation of lentiviruses expressing *Cre* recombinase (DuPage et al., 2011). KP mice crossed to *Rosa26*^LSL-Cas9-GFP-Csy4^ (Ng et al., 2020) and the *Rosa26*^LSL-tdTomato^ were used for CRISPR/Cas9-mediated gene activation of Cxcl10. Mice were between 8 and 14 weeks of age at the time of lentiviral infection. Males and females were used equally across all experimental arms. All studies were performed under an animal protocol approved by the Massachusetts Institute of Technology (MIT) Committee on Animal Care. Mice were assessed for morbidity according to guidelines set by the MIT Division of Comparative Medicine and were humanely sacrificed prior to natural expiration. Information about each mouse experiment is provided in **Table S4**.

### Lentiviral Tumor Induction

To initiate lung tumors, KP mice were injected intratracheally (i.t.) with 2.5 x 10^4^ PFU of lentivirus containing *Cre* recombinase and model neoantigens *as* previously described (DuPage et al., 2009, 2011). Details of the lentivirus production can be found below. Mice were randomized post-infection for immunotherapy trials.

### Lentiviral Constructs

Lentiviral constructs containing *Cre* recombinase with or without LucOS antigens (Lenti-Cre and Lenti-LucOS) were previously described (DuPage et al., 2011). The Lenti-Cre design was modified by Gibson cloning to create Lenti-SAM-Cre for CRISPR/Cas9-mediated gene activation. A U6 promoter and an activator guide RNA cloning cassette were added upstream and inverted from the Pgk promoter driving *Cre*. The cloning cassette contains BsmBI restriction sites for the addition of a 15-nucleotide “dead” guide RNA (dRNA) to mediate gene activation rather than cutting by catalytically active Cas9 (Dahlman et al., 2015). The cassette appends the dRNA with stem-loops containing MS2-binding aptamers as previously described (Konermann et al., 2015). “SAM” transcriptional activation components from p65 (NFkB) and Hsf1 were fused with the MS2 RNA binding protein (Dahlman et al., 2015; Konermann et al., 2015) and cloned in tandem with *Cre*, separated by a P2A self-cleaving peptide. For *in vitro* validation of dRNA activity, Lenti-SAM-Cre was modified to replace *Cre* with a Puromycin selection gene (Lenti-SAM-Puro).

### Cxcl10 Dead Guide RNA Screening

Short guide RNA (sgRNA) sequences targeting the promoter region of *Cxcl10* (up to 200 nucleotides upstream of the TSS) were selected using the Feng Zhang lab (Broad Institute of MIT and Harvard) online SAM Cas9 activator design tool (no longer operational). The 20 nucleotide sgRNA sequences were shortened to 15-nucleotide dead RNAs (dRNAs) to recruit Cas9 to the promoter region but prevent DNA cleavage by Cas9. The first nucleotide was amended to a G if it did not occur naturally to optimize expression from the U6 promoter. The dRNAs were screened for their relative ability to activate *Cxcl10* expression in the 1233 KP lung adenocarcinoma cell line. Briefly, oligonucleotides were generated with BsmBI restriction site overhangs (see Key Resources Table) and annealed to create the double-stranded dRNAs for cloning into Lenti-SAM-Puro. 293FS* viral packaging cells were transfected in a 6- well plate format with the dRNA-containing Lenti-SAM-Puro constructs (1.5 µg) and psPAX2 (0.75 µg) and VSV-G (0.25 µg) helper plasmids to generate lentivirus. The lentiviral supernatant was collected through a 0.45 µm filter 48 hrs post-transfection and added 1:1 to 1233 KP Cas9 cells plated at 25,000 cells/well the day before. Polybrene was added to improve transduction efficiency at 4 µg/ml. Puromycin was added 48 hrs later to select for cells expressing the construct. Cells were expanded (under Puromycin selection) and plated in triplicate in 12-well plates at 200,000 cells/well to generate supernatant containing secreted Cxcl10. The supernatant was collected 72 hrs later and Cxcl10 protein was quantified using a Cxcl10 ELISA (R&D systems) according to the manufacturer’s protocol. The dRNA that resulted in the greatest production of Cxcl10 (GACAAGCAATGCCCT) was cloned into Lenti- SAM-Cre and used to generate large-scale lentivirus for *in vivo* studies. A non-targeting dRNA shortened from an sgRNA targeting tdTomato (CGAGTTCGAGATCGA; (Sánchez-Rivera et al., 2014) was used a negative control. dRNA sequences and oligonucleotides are listed in the key resources table.

### Lentivirus Production for *In Vivo* Instillation

Lentivirus was produced by transfection of 293FS* viral packaging cells in 15 cm plates with lentiviral constructs (10 µg), VSV-G (2.5 µg) and psPAX2 (7.5 µg) viral packaging plasmids, and Mirus TransIT LT1 (MirusBio; 60 µl). Lentiviral supernatant was harvested, passed through a 0.45 um filter, and concentrated by ultracentrifugation at 25,000 rpm for 2 hrs at 4°C 48- and 72-hrs post-transfection. Viral titers were determined by measuring *Cre* activation of GFP expression in GreenGo 3TZ cells as previously described (Sánchez-Rivera et al., 2014).

### Anti-PD-1/Anti-CTLA-4 Therapy

KP LucOS mice were treated for one week starting at 8 wks post-tumor initiation with InvivomAb anti-PD1 (29F.1A12; BioXCell) and InvivomAb anti-CTLA4 (9H10; BioXCell) or isotype controls (Rag IgG2a, 2A3; Syrian Hamster, polyclonal; BioXCell). Mice received 200 µg of each antibody i.p. at day 0, followed by 200 µg anti-PD-1 and 100 µg anti-CTLA-4 (or isotype controls at the same concentrations) on days 3 and 6. Mice were sacrificed for endpoint analysis on day 7.

### Neoantigen Vaccination

KP LucOS mice were vaccinated s.c. at the tail-base with 30 amino acid long peptides containing SIINFEKL and SIYRYYGL (10 nmol; New England Peptide) and cyclic-di-GMP adjuvant (0.25 mg/ml; Invitrogen) at 6 wks post-tumor initiation. An equivalent booster dose was given 2 wks later, and the mice were sacrificed at 9 wks post-tumor initiation for endpoint analysis. All doses were delivered in two 50 µL boluses and control mice received PBS. The long peptide sequences used were: SMLVLLPDEVSGLEQLESIINFEKLTEWTS and GRCVGSEQLESIYRYYGLLLKERSEQKLIS (New England Peptide).

### Mouse Lung Tissue Processing for Histology and H&E Staining

Tumor-bearing lung lobes were collected into 4% paraformaldehyde in PBS and incubated overnight with shaking at 4°C. Tissue was transferred into 70% ethanol and subsequently paraffin embedded and sectioned (4 µm) onto Fisherbrand Superfrost Plus Microscope Slides (ThermoFisher Scientific). After drying, slides for RNAScope™ were stored at 4°C until use. Hematoxylin and eosin (H&E) stain was performed with a standard method by the Hope Babette Tang Histology Facility at the Koch Institute at MIT.

### Tissue-Based Cyclic Immunofluorescence (t-CyCIF) Staining and Imaging

FFPE sections were prepared and stained with a 24-plex antibody panel according to the previously described t-CyCIF protocols (Burger et al., 2021; Gaglia et al., 2022; Lin et al., 2018) (see **Table S1**). This CyCIF panel has been validated across many different sample types in accordance with standards defined by our group (Du et al., 2019).

#### Baking and Dewaxing

To prepare samples for antibody staining, slides were automatically baked at 60°C for 30 min, dewaxed at 72°C in BOND Dewax Solution, and antigen retrieval was performed at 100°C for 20 min in BOND Epitope Retrieval Solution 2 (ER2) by the Leica Bond RX machine.

#### Pre-Staining Background Reduction

After slides were baked and dewaxed, they were photobleached by immersing them in bleaching solution (4.5% H2O2, 20 mM NaOH in PBS) with LED light exposure for 2 x 45 min to reduce autofluorescence.

To mitigate non-specific antibody binding, slides were washed for 3 x 5 min with 1X PBS and then incubated overnight with secondary antibodies (anti-rat, anti-mouse, and anti-rabbit) diluted in 150 μL of Odyssey Blocking Buffer (1:1000) at 4°C in the dark. Slides were subsequently washed 3x with 1X PBS before photobleaching them again for 2 x 45 min.

#### Antibody Staining, Slide Mounting, and Imaging

For each round of t-CyCIF, samples were incubated overnight at 4°C in the dark with Hoechst 33342 (Dilution: 1:10,000; Thermo Fisher Scientific, cat# 62249) for nuclear staining along with either primary conjugated antibodies or primary unconjugated antibodies diluted (see **Table S1** for antibody information) in 150 μL of Odyssey Blocking Buffer (LI-Cor, Cat# P/N 927–40003). Incubation with primary unconjugated antibodies was followed by secondary antibody incubation at room temperature for 2 hrs in the dark.

Post staining, slides were washed for 3 x 5 min, mounted with 24 x 50 mm coverslips using 200 μL of 70% glycerol, and then dried. Once coverslipped, slides were manually imaged on the IN Cell Analyzer 6000 or automatically on the RareCyte Cytefinder II HT using the following channels: UV, cy3, cy5, and cy7 (Binning: 1 x 1; Objective: 20×; Numerical Aperture: 0.75; Resolution: 0.325 μm/pixel). Image exposures were optimized for each channel to avoid signal saturation and kept constant for each sample

To demount, slides were placed in containers of 1X PBS and heated in a water bath for 1 hr. Before additional antibody staining, slides are photobleached for 2 x 45 min to deactivate the fluorophores and washed 3 x 5 min in 1X PBS.

#### RNA In Situ Hybridization

RNAScope™ was performed as per manufacture’s suggested protocol (Advanced Cell Diagnostics, Inc.) using the LS Multiplex Reagent Kit (cat# 322800) and probes RNAscope® 2.5 LS Probe- Mm-Cxcl9 (cat #: 489348) and RNAscope® 2.5 LS Probe- Mm-Cxcl10-C3 (cat #: 408928-C3).

### Quantification and Statistical Analysis

Information on the sample size and the statistics are included in the figure legends. Statistical tests used are Pearson correlation, two-sided t-test, and non-parametric Kolmogorov–Smirnov (KS) two-sided test as specified in the figure legends and are performed with MATLAB built-in functions. Significance was defined as a p-value of less than 0.05.

### Image Processing and Single-Cell Quantification

The image processing of both plate-based and tissue cyclic immunofluorescence was organized in the following steps, each of which is described in detail below:

- the software ASHLAR is used to stitch, register, and correct for image acquisition artifacts (using the BaSiC algorithm). The output of ASHLAR is a single pyramid ome.tiff file for each region imaged;
- the ome.tiff file is re-cut into tiles (typically 5000 x 5000 pixels) containing only the highest resolution image for all channels. One random cropped image (250 x 250 pixels) per tile is outputted for segmentation training (using Fiji);
- the ilastik software is trained on the cropped images to label, nuclear, cytoplasmic, and background areas. The output of the Ilastik processing is a 3-color RGB image with label probabilities;
- the RBG probability images are thresholded and watershed in MATLAB to segment the nuclear area. The cytoplasmic measurements are derived by dilating the nuclear mask;
- single-cell measurements are extracted for each channel (cell pixel median and mean for both nuclear and cytoplasmic area) as well as morphological measurements of area, solidity, and cell coordinates location.

#### BaSiC

The BaSiC ImageJ plugin tool was used to perform background and shading correction of the original images (Peng et al., 2017). The BaSiC algorithm calculates the flatfield, the change in effective illumination across an image, and the darkfield, which captures the camera offset and thermal noise. The dark field correction image is subtracted from the original image, and the result is divided by the flatfield image correction to obtain the final image.

#### ASHLAR

Alignment by Simultaneous Harmonization of Layer/Adjacency Registration (ASHLAR) is used to stitch together image tiles and register image tiles in subsequent layers to those in the first layer (Muhlich et al., 2021). For the first image layer, neighboring image tiles are aligned to one another via a phase correlation algorithm that corrected for local state positioning error. A similar method is applied for subsequent layers to align tiles to their corresponding tile in the first layer. ASHLAR outputs an OME-TIFF file containing a multi-channel mosaic of the full image across all imaging cycles. Full codes available at: https://github.com/labsyspharm/ashlar.

#### ilastik

ilastik is a machine learning based bioimage analysis tool that is used to obtain nuclear and cytoplasmic segmentation masks from OME-TIFF files (Berg et al., 2019). For increased processing speed, randomly selected 250 x 250 pixel regions from the original OME-TIFF are used as training data. ilastik’s interactive user interface allows the user to provide training annotations on the cropped regions. Users are presented with a subset of the channels stacked images and label pixels as either nuclear area, cytoplasmic area, or background area. The annotations are used to train non-linear classifiers that are applied to the entire image to obtain probability masks describing the probabilities of each pixel belonging to the nuclear, cytoplasmic, or background area. A MATLAB (version 2018a) script uses these masks to construct binary masks for nuclear and cytoplasmic area.

#### Single Cell Segmentation and Quantification

Using ilastik’s Pixel Classification workflow, a random forest classifier is trained for each experimental dataset based on manual annotations of nuclear, cytoplasmic, and background regions within the CroppedData. Batch processing is subsequently performed by the classifier on the FullStacks, generating .tif probability maps for nuclei, background, and cytoplasm.

Cell nuclei are segmented through thresholding maps based on nuclear, cytoplasm, and background probabilities and performing water shedding on them using MATLAB. Cytoplasmic segmentation masks are produced by dilating nuclear segmentation masks radially by 3 pixels and then excluding the segmented nuclear area.

Median nuclear and cytoplasmic marker expression, centroid coordinates, area (nuclear and cytoplasmic), and solidity are quantified for each segmented cell using MATLAB’s regionprops function and outputted as a single “Results.mat” file for each FFPE slide. All MATLAB scripts used for segmentation and quantification can be found here: https://github.com/santagatalab.

### Data analysis workflow

The data analysis is divided in a set of pre-processing steps in which data from different tissues is i) log2-transformed and aggregated together, ii) filtered for image analysis errors, and iii) normalized on a channel-by-channel basis across the entire data from a single experiment. All the steps are performed in MATLAB.

#### Data aggregation

The image processing workflow outputs one ome.tiff image and one data file (.mat) for each tissue area imaged. The data matrices from each .mat file are concatenated into a single matrix for each metric measured (median/mean, nuclear/cytoplasmic) into a single structure (“AggrResults”). The morphological data (i.e., area, solidity, and centroid coordinates) is concatenated into a single structure (“MorpResults”), which also contains the indexing vector to keep track of the tissue of origin within the dataset.

#### Data filtering

Single cells are filtered to identify and potentially exclude from subsequent analysis errors in segmentation and cells lost through the rounds of imaging. Two types of criteria are used to filter cells: morphological criteria based on cell object segmented area, which are applied to all the rounds for the cell object, and DAPI-based criteria which are applied to the DAPI measurement for each imaging round. The latter corrects for cell loss during cycling and computational misalignment, which are both round specific.

Morphological filtering criteria are:

- nuclear area within a user-input range;
- cytoplasmic area within a user-input range;
- nuclear object solidity above a user-input threshold.

DAPI-based criteria are:

- nuclear DAPI measurement above a user-input threshold;
- ratio between nuclear and cytoplasmic DAPI measurement above a user-input threshold;

The filter information for the criteria is allocated to a logical (0-1) structure ‘Filter’, which is used to select the cells to analyze in the further analysis by indexing. The threshold selection is dataset dependent and is performed by data inspection. The values used in each dataset are available with the codes used for data analysis in the Synapse.org repository syn30715952.

#### Data normalization

Each channel distribution is normalized by probability density function (pdf) centering and rescaling. The aim is to center the distribution of the log2 fluorescent signal at 0 and rescale the width of the distribution to be able to compare across channels. The data is first log-transformed (base 2). The standard normalization is performed using a 2-component Gaussian mixture model, each model capturing the negative and the positive cell population. If the 2- component model fails to approximate the channel distribution, two other strategies are attempted: i) a 3-component model is used assuming the components with the two highest means are the negative and positive distribution (i.e., discarding the lowest component) or ii) the user selects a percentage ‘x’ of assumed positive cells and a single Gaussian distribution fit is performed on the remainder of the data to capture the negative distribution. The single Gaussian fit is then used as the lower component in a 2-component model to estimate the distribution of the positive population. The strategy chosen for each channel in each dataset is available in the code section of the Synapse.org repository syn30715952.

The “add_coeff” is defined as the intersection of the negative and positive distributions. The “mult_coeff” is defined as the difference between the mean of the negative and positive distributions. The full distribution is normalized by subtracting the add_coeff and dividing by the mult_coeff. The normalization is performed on the nuclear and cytoplasmic single-cell, single-channel distributions individually.

The data preprocessing workflow is performed on all datasets. The individual analyses used in the paper are performed only in selected datasets as follows.

#### Cell type classification

Cell type classification is performed hierarchically on the filtered, normalized expression data. Each cell is evaluated based on marker expression and then assigned to cell types in a layered fashion according to the dendrogram schematic in **Figure S1C**, with each successive layer being more specific than the previous one. A cell is considered to be positive for a marker if its median expression is above 0. Cell types are defined in the dendrogram by the presence or exclusion of multiple markers using “&&” and “||” operators representing “AND” and “OR” logic respectively. If multiple marker conditions must be met to assign a cell type, these marker conditions are grouped using parentheses. If a cell is “positive” for two markers that are expected to be mutually exclusive, the marker that is expressed at a higher value takes precedence as long as the difference in expression surpasses a user-defined threshold.

#### Multimodal Data Integration

H&E, RNAScope™ and CyCIF images are rescale and registered using the open-source software elastix (Klein et al., 2010) using non-shearing global transformation. The CyCIF images are used as the fixed images in elastix. To integrate the CyCIF and histological data, H&Es are annotated for tumors, blood vessels, and airways by a trained pathologist. The elastix registration is used to overlay the pathology annotation onto the CyCIF single cell coordinates and then to calculate the distance from tumor boundaries and blood vessels.

#### RNAscope foci detection

Custom spot detection scripts (https://github.com/Yu-AnChen/wsi-fish) are used to identify RNAScope™ foci and quantify their intensity. Each RNAScope™ dot is assigned as belonging to the closest cell based on the segmented area. A cell is considered Cxcl positive if it is assigned at least two RNA foci and if the cumulative RNAScope™ dot intensity of all the dots assigned to the cell exceed a preset threshold (based on the positive tail of the single cell distribution).

#### Lymphonet definition

The single cell centroids are tessellated using the Delaunay Triangulation using a custom script in MATLAB (https://github.com/santagatalab) to obtain a 2D graph, setting a maximum edge length of 16.25 microns (50 pixels). Using conventional graph operations, the graph edges are then filtered to include only connection between lymphocytes (Lv3 of cell type dendogram), after which connected subgraphs of length greater than 5 are than defined as “lymphonets”.

#### Palantir algorithm and CD8 T cell state definition

The algorithm Palantir (Setty et al., 2019) was adapted to CyCIF data by bypassing the initial dimensionality reduction applied to single-cell RNA-seq data and using the CyCIF channel information as the dimensionality reduction output. The Python Jupiter Notebooks used to run the Palantir analyses can be found at https://github.com/santagatalab. The CD8 T-cell phenotypic states S1-S3 and T1-T3 were obtained using a flow cytometry manual gating approach combining Palantir point density and marker intensity. The gating was performed in MATLAB using the “Flow Cytometry GUI for Matlab” by Nitai Steinberg (2022) available at https://www.mathworks.com/matlabcentral/fileexchange/38080-flow-cytometry-gui-for-matlab, MATLAB Central File Exchange.

#### Spatial Correlation Analysis

Spatial correlations Cxy(r) were computed as the Pearson correlation between a cell of group X and its kth nearest neighbor of group Y, for their respective variables x and y. A value of Cxy (r) was computed for each k up to 100, and a distance r was assigned to each k as the average distance between kth nearest neighbors. More detail can be found in Gaglia et al., 2022.

### Visinity - Visual Spatial Neighborhood Analysis

To visually explore the spatial neighborhoods within these data, we use the *Visinity (*Warchol et al., 2022), a scalable system for visual analysis in whole-slide multiplexed tissue imaging data. This system supports the analysis of recurrent cellular spatial neighborhoods across cohorts of specimens. Visinity is an open-source project (https://github.com/labsyspharm/visinity), with a JavaScript client for browser-based visualization and a Python server for efficient and scalable backend computation.

#### Quantifying Cellular Neighborhoods

Visinity quantifies the spatial neighborhood for each cell in terms of the types of cells that surround it (for Visinity the information contained in level 4 (Lv4) was used as the cell type information). More specifically, this process is as follows:

A ball-tree index structure is constructed using nuclei centroids of each segmented cell in a specimen, which allows for *O(n + k)* range queries, where *n* is the number of cells and *k* is the number of points within this range. We use the scikit-learn (Pedregosa et al., 2011) implementation of this data structure.

With the ball-tree, we identify neighboring cells within a 50 μm radius of each cell.

We create feature vectors representing the neighborhood of each cell. Vectors are 1 x *n*, where *n* is the number of cell types. Columns in this vector correspond to the presence of a specific cell type. We linearly weight each cell in a neighborhood by its distance from the center so that cells just at the edge of the neighborhood radius contribute the least and sum these weights by cell type.

We repeat this process for every cell across all specimens, L1 normalizing the vectors. Each vector, which represents the neighborhood of an individual cell, is a row in a matrix representing all cells across all specimens.

We create a 2D embedding of this matrix using UMAP (McInnes et al., 2018) with the parameters *n_neighbors = 50, min_dist = 0.1*. Points close to each other in this embedding space represent cells with similar spatial neighborhoods

We display this embedding as an interactive scatterplot. Selecting regions in this embedding highlights the corresponding cells within the tissue image and we visualize the cell types that compose the selected neighborhood with a parallel coordinates plot.

Visinity supports both confirmatory and exploratory analysis, allowing users to detect spatial neighborhood patterns in a semi-automated manner and visually query across specimens for specific cellular neighborhoods. This workflow and the system as a whole are described in detail in Warchol et al., 2022.

## RESOURCE AVAILABILITY

### Data and code availability

Datasets generated and the corresponding analysis used in all figures will be made available in the Synapse.org repository (www.synapse.org/#!Synapse:syn30715952/wiki/617734). Multiplexed images of a mouse lung specimen (KP LucOS can be viewed in Minerva Story (Hoffer et al., 2020; Rashid et al., 2022) an interpretive guide for interacting with multiplexed tissue imaging data https://tinyurl.com/mouseprofiling. Imaging data is available from the corresponding authors on request. MATLAB codes used to perform the t-CyCIF processing and analysis are available on GitHub: https://github.com/santagatalab. Code for analysis of recurrent cellular spatial neighborhoods using the Visinity algorithm (Warchol et al., 2022) is available on GitHub: https://github.com/labsyspharm/visinity.

## KEY RESOURCES TABLE

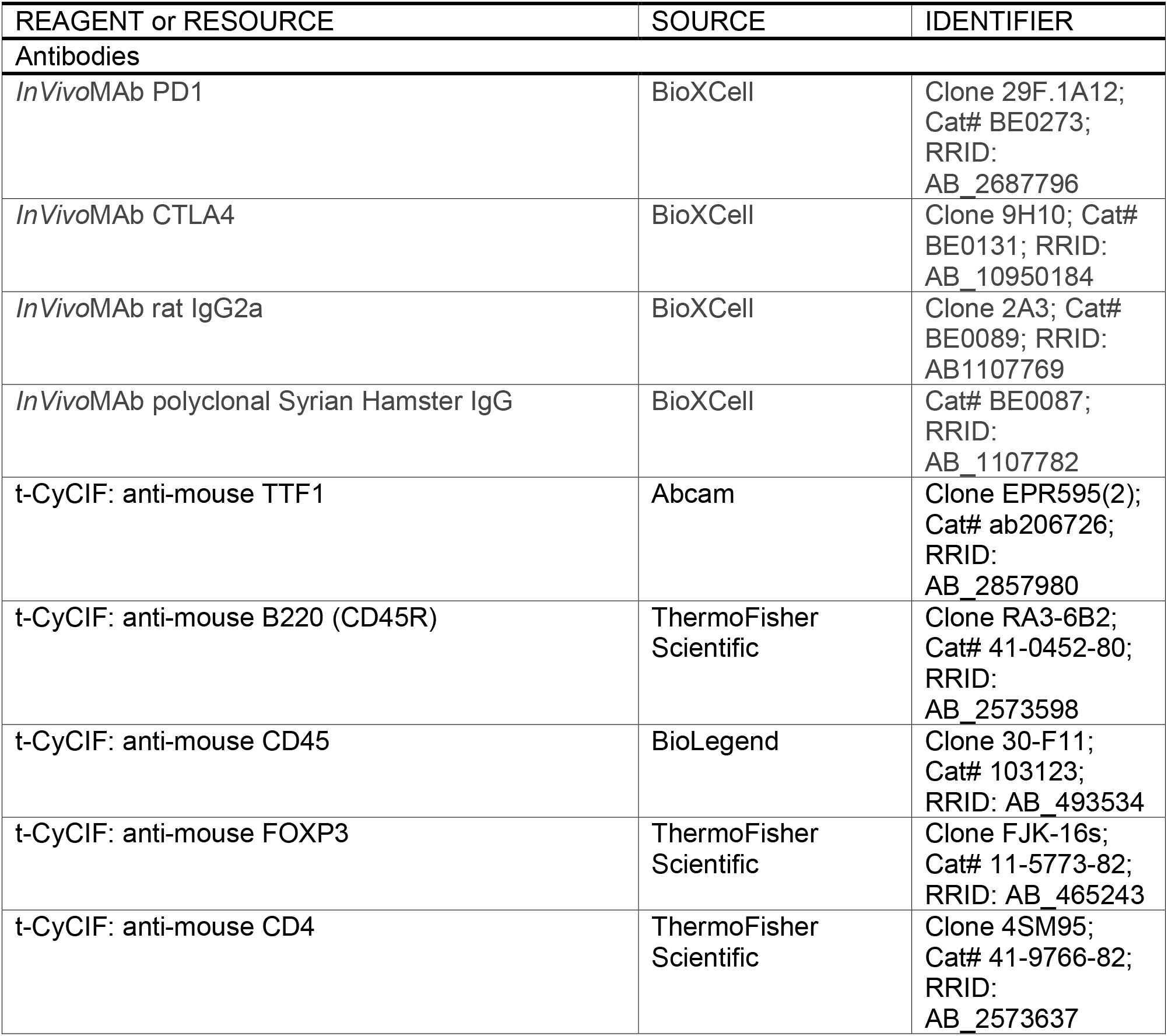
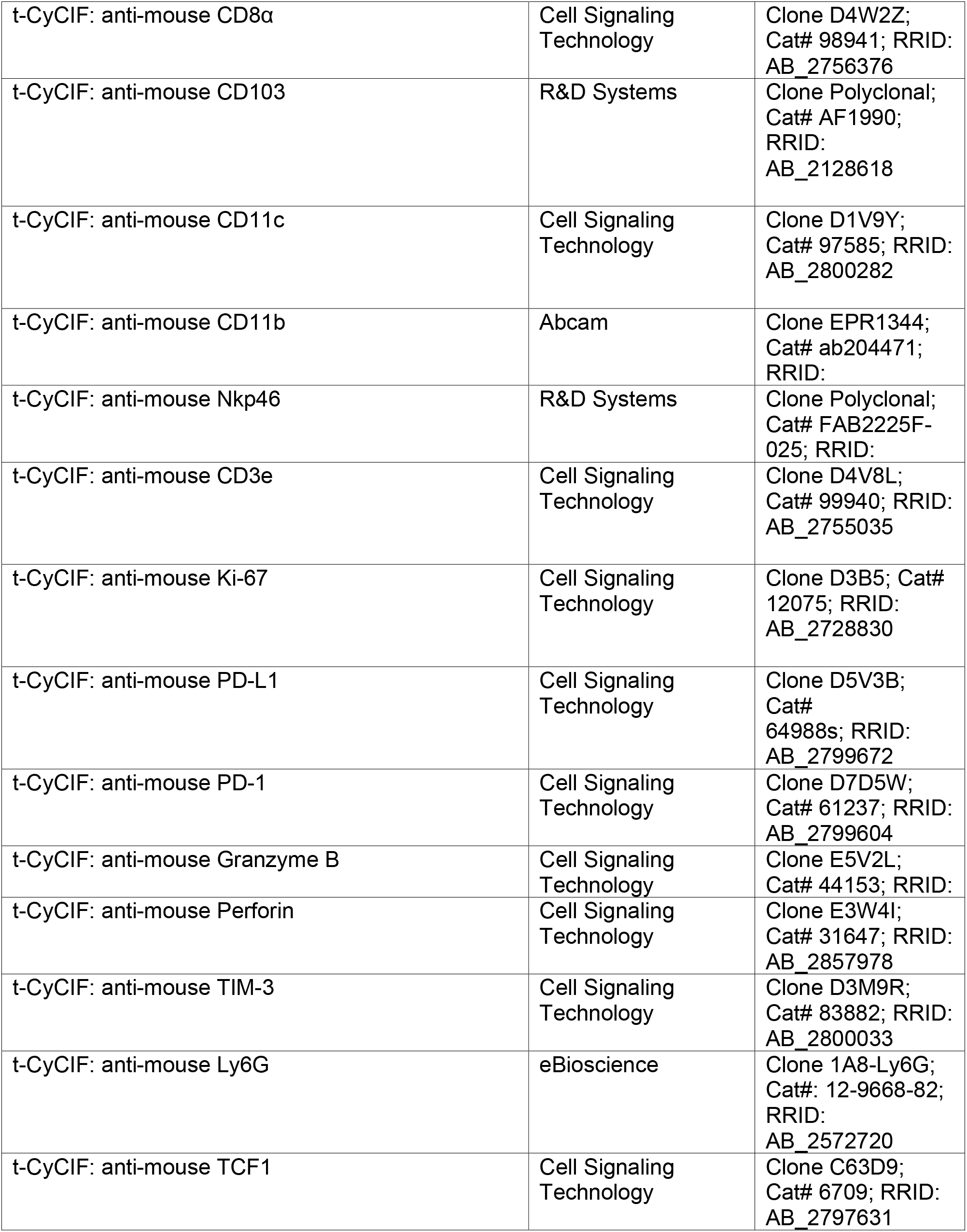
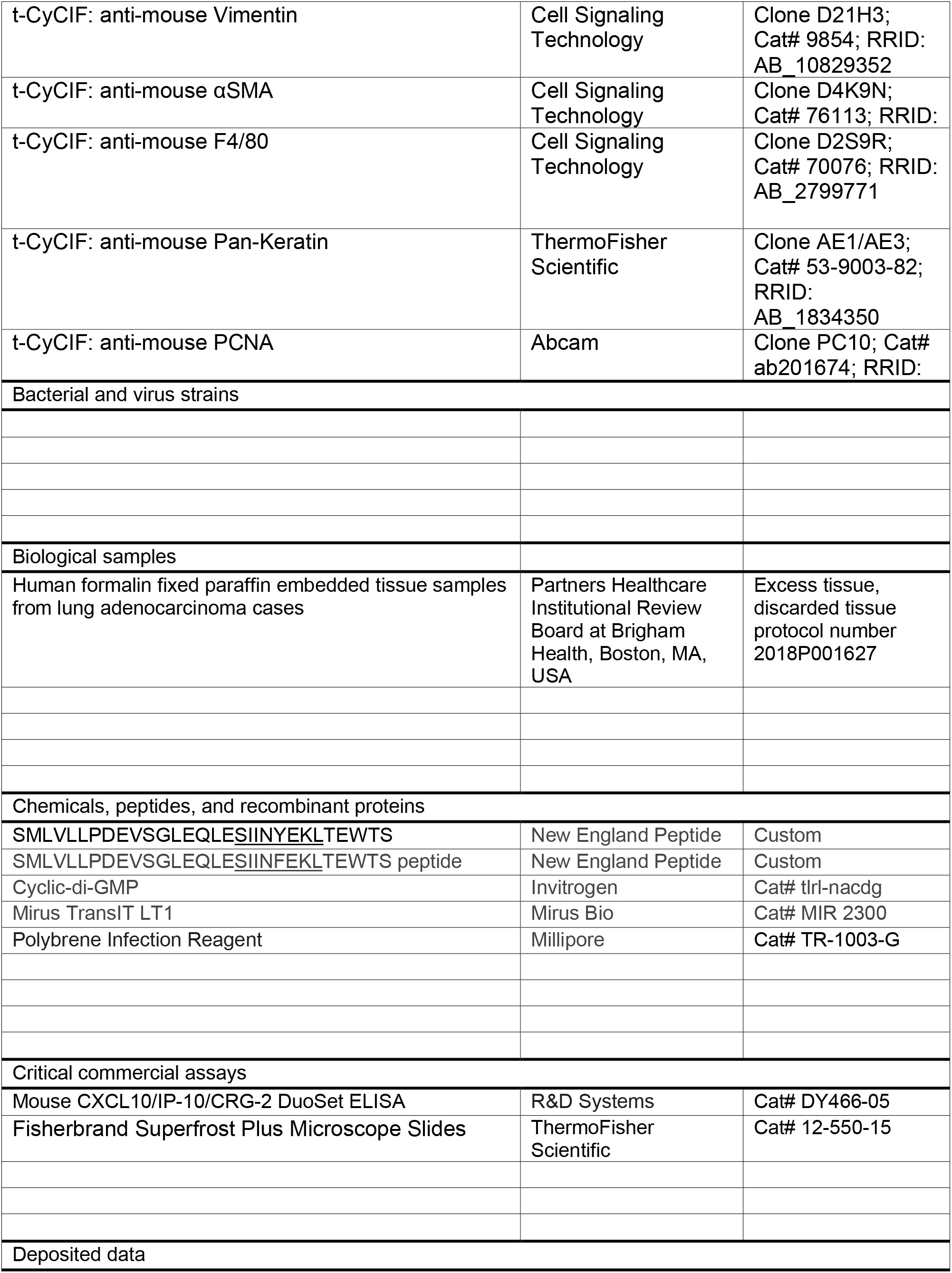
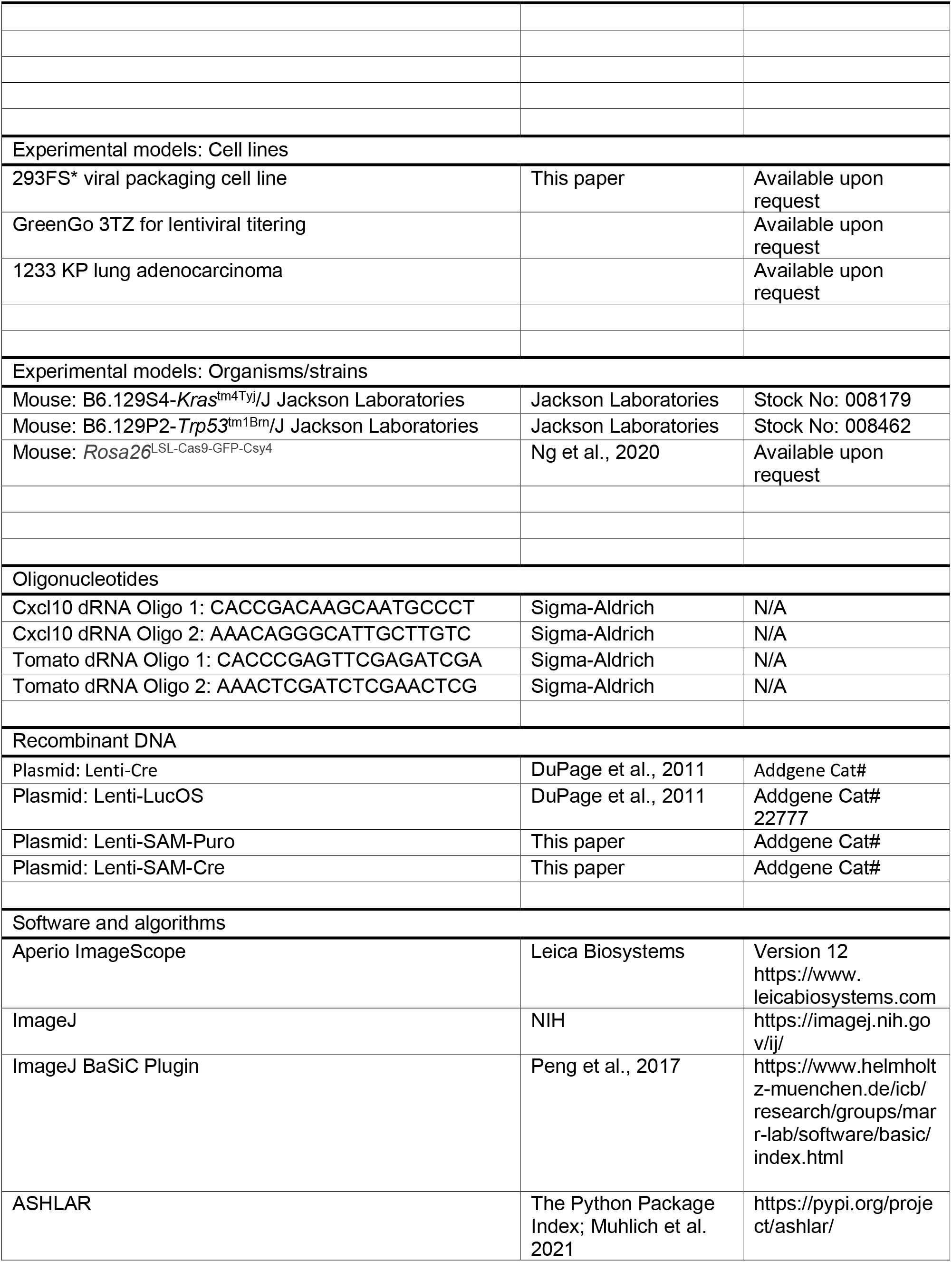
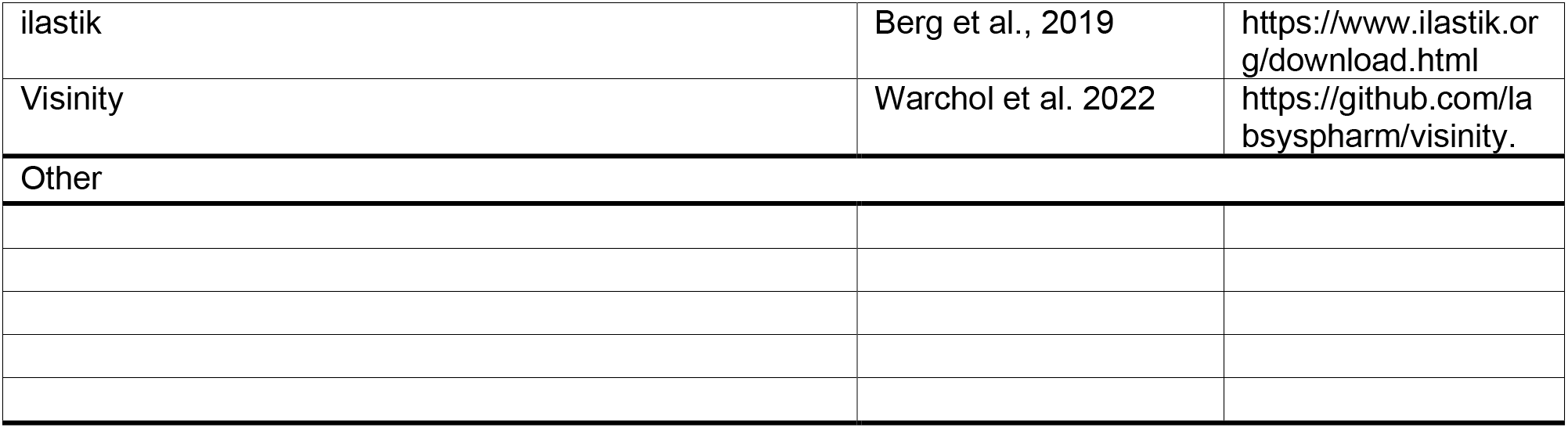

---

**Figure S1.**
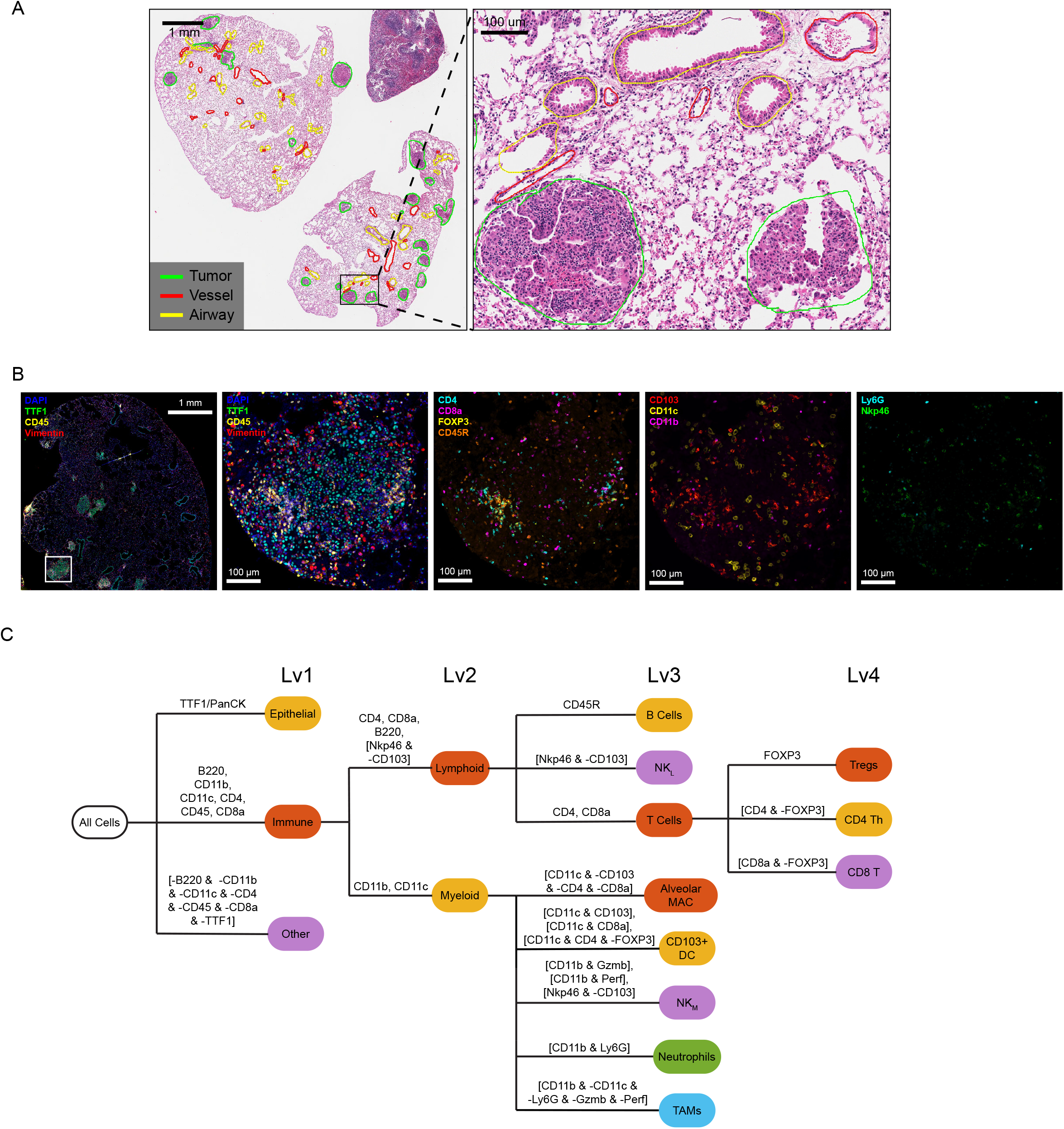
Multiplexed tissue imaging and cell type calling in the KP genetically engineered mouse model of cancer. (A) Representative image of H&E-stained section with pathology annotations indicated. (B) Representative multiplexed CyCIF images (whole lung lobe and single tumor inset) of tumor and immune markers from whole FFPE sections of KP LucOS tumor-bearing lung. (C) Cell type calling dendrogram for CyCIF image analysis; first immune, epithelial, and ‘other’ cell types were identified (Level 1, Lv1; shown in heat map of all cells), and then the immune cells were further clustered into lymphoid cells and myeloid cells (Level 2; Lv 2) and immune cell subtypes (Level 3 and Level 4; Lv3, Lv4: Treg, CD4 Th, CD8 Tc, B cells, NK cells (lymphoid marker-defined, ‘NK-L’), alveolar macrophages, dendritic cells, NK cells (myeloid marker-defined, ‘NK-M’), neutrophils, and tumor associated macrophages (CD11b+CD11c-).

**Figure S2.**
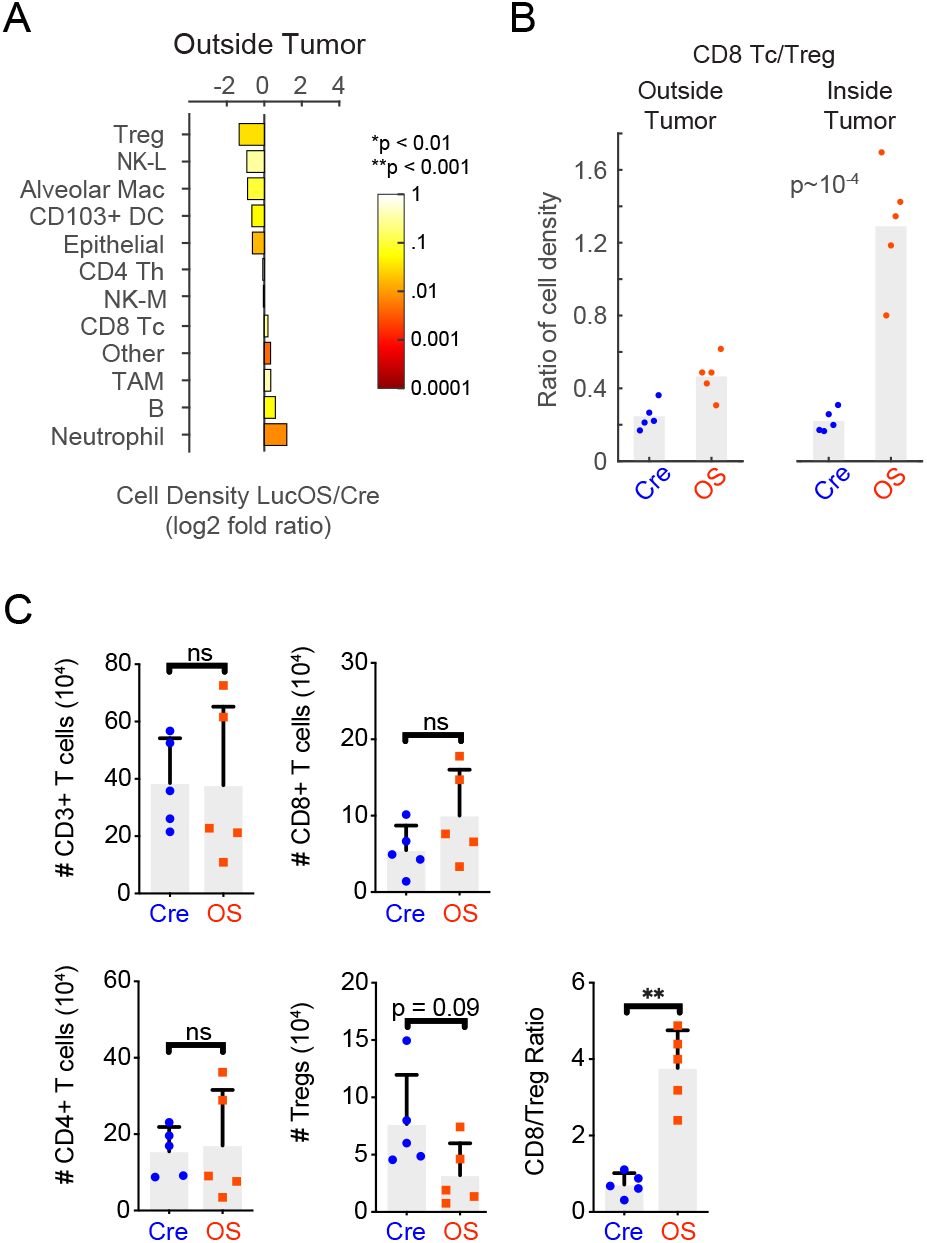
Spatial analysis of immune cell-type composition in KP LucOS versus Cre tumor-bearing lung. (A) Log2 fold ratio of cell-type densities between LucOS and Cre in areas outside tumor (n = 5 mice per group, color represents two tailed t-test p-value). (B) Ratio of CD8 Tc to Treg cell density measurements outside and inside of annotated tumor areas in LucOS versus Cre mice (n = 5 mice per group, bar = mean). (C) T-cell numbers in LucOS versus Cre mice calculated by flow cytometric analysis of dissociated tumor-bearing lung tissue (n = 5 mice per group, bar = mean).

**Figure S3.**
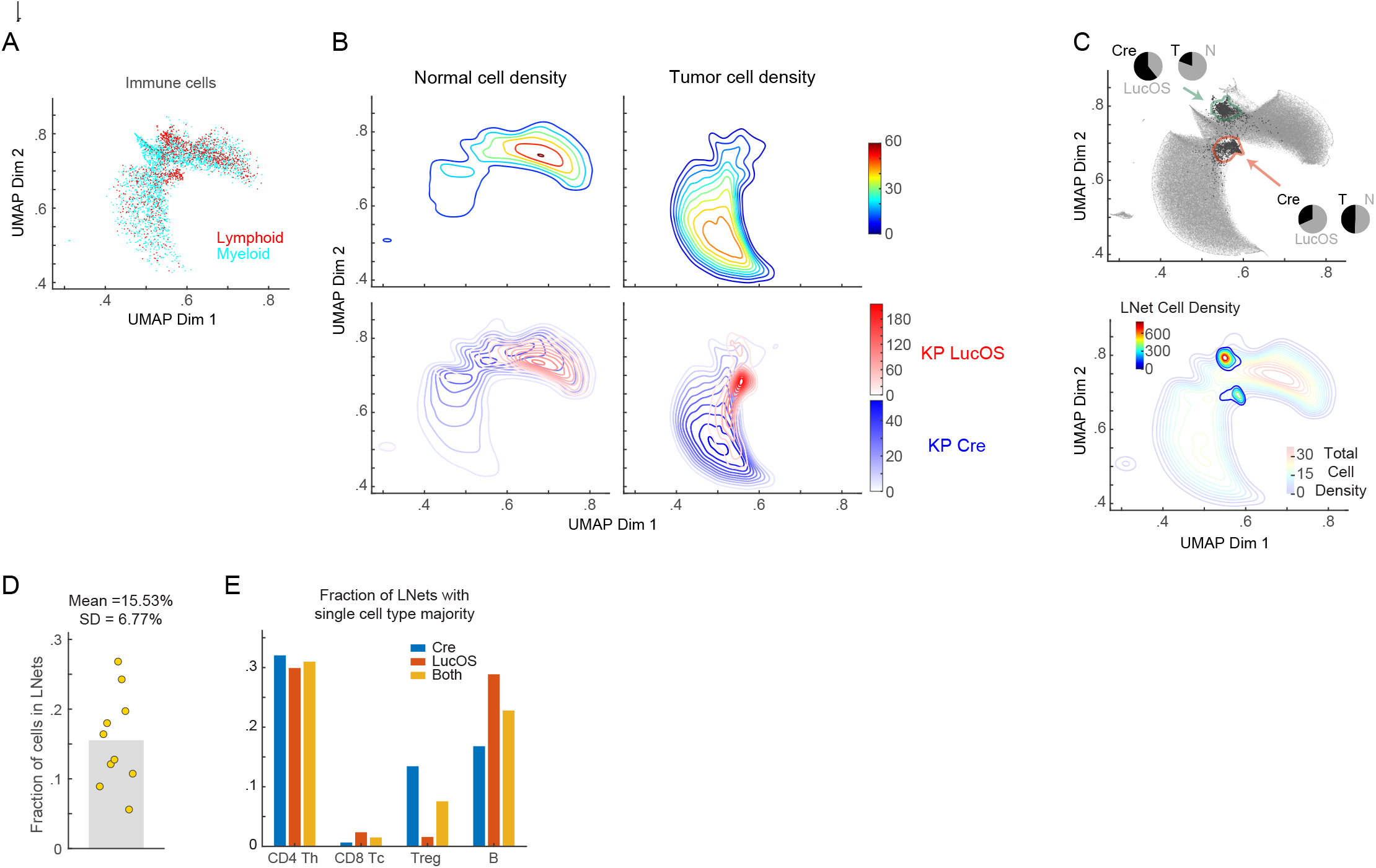
Characterization of lymphocyte networks. (A) Neighborhood embedding generated by the Visinity algorithm displaying all cells. (B) Density plots of normal and tumor cells in Visinity embedding of KP LucOS and KP Cre lung tissue. (C) Visinity plots, black dots are cells in lymphonets. Arrows indicate Visinity cluster enriched in lymphonets, and pie charts summarize the composition and fraction of each cluster derived from LucOS and Cre and from tumor and normal tissues. (D) Plot of fraction of cells in lymphonets for LucOS and Cre together (n = 10 mice, 5 per group, bar = mean). (E) Fraction of lymphonets with indicated single-cell type majority for LucOS, Cre, and combined (n = 5 mice per group).

**Figure S4.**
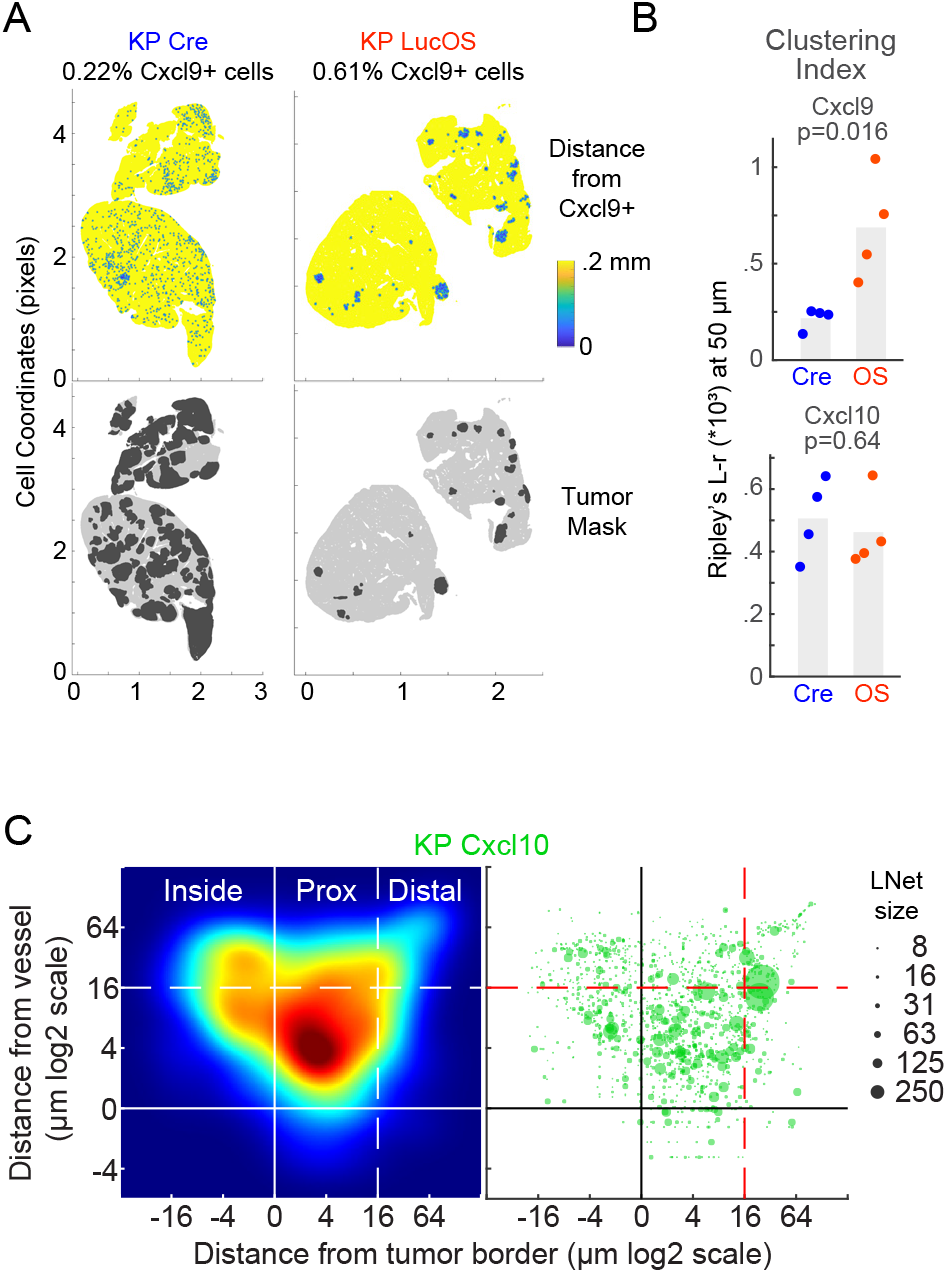
Spatial analysis of *Cxcl10* overexpression on lymphonets. (A) Map of all cells colored by distance to nearest *Cxcl9* mRNA positive cell measured by RNAScope™ *in situ* hybridization in KP Cre and KP LucOS lung tissue. (B) Spatial autocorrelation of *Cxcl9* and *Cxcl10* mRNA-expressing cells using Ripley’s L function (‘Ripley’s clustering index’) in KP Cre and KP LucOS mice (n = 4 mice per group, bar = mean). (C) Left, density plots of lymphonets by distance from closest blood vessel (y-axis) and tumor (x-axis) for KP *Cxcl10* cohort. Right, scatter plot lymphonets used to generate density plot (dot size represents the lymphonet size (n = 5 mice per group).

**Figure S5.**
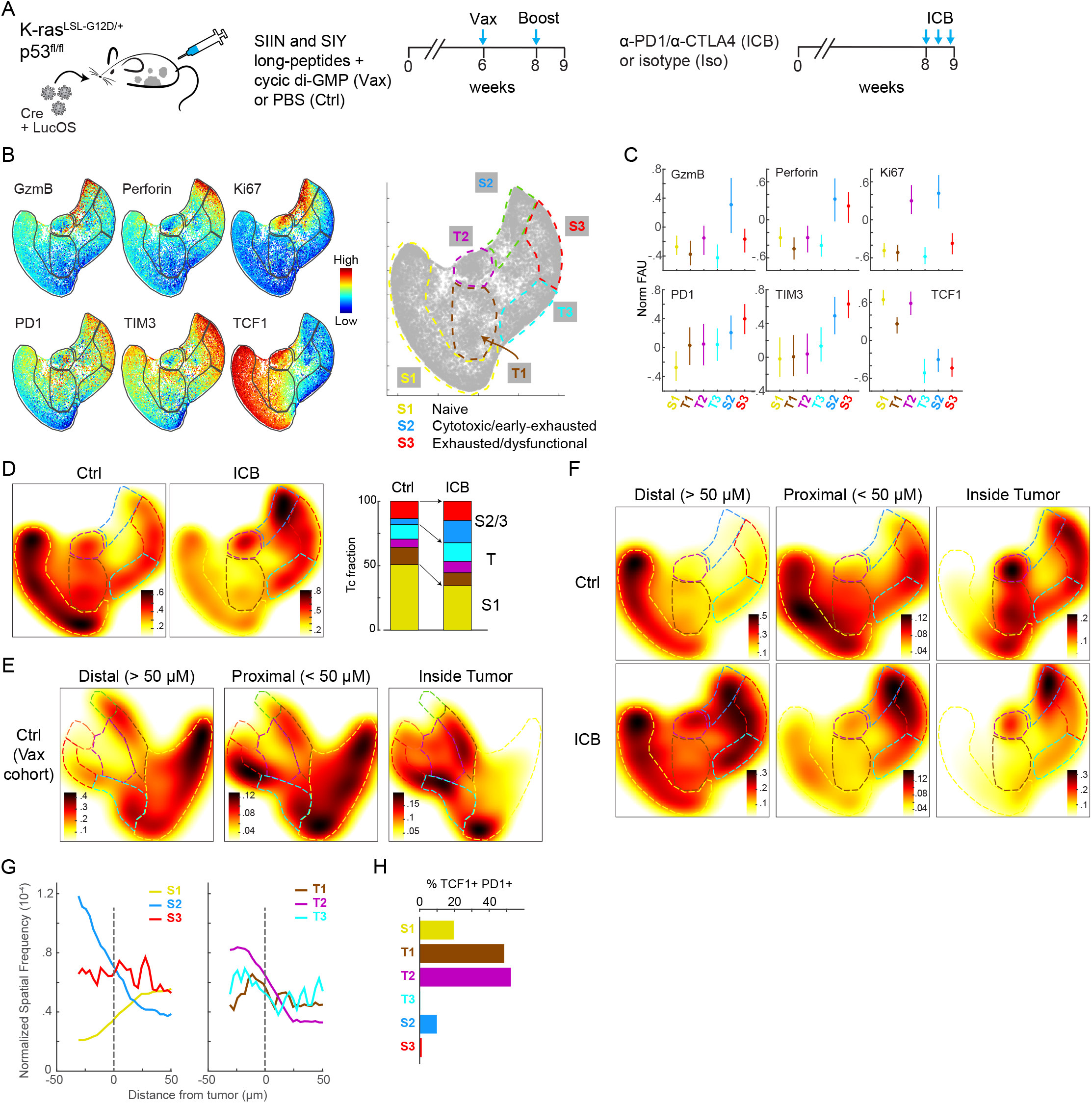
Multiparametric analysis of Tc functional states after anti-PD1/anti-CTLA4 immune checkpoint blockade (ICB) (A) Schematic of treatment of KP LucOS mice with a SIIN and SIY long-peptide vaccine or anti-PD-1/anti-CTLA4 immune checkpoint blockade therapy. (B) Palantir projection of Tc populations in KP LucOS mice treated with anti-PD1/anti-CTLA4 immune checkpoint blockade (ICB) or isotype control antibodies (Ctrl) (n = 10^4^ cells sampled from n = 6 per treatment, see S5A for treatment schematic). The expression levels of the indicated markers are mapped to color. Tc states defined by multiparameter measurements are indicated at the extremes of the representation (S1, S2, and S3) connected by transitional phenotypes (T1-T3) shown in the schematic to the right. (C) Plot of the normalized fluorescence units for each of the markers in the indicated Tc cell states and transitions. (D) Heat map of Tc cell densities in Palantir projections for Ctrl and ICB groups (n = 10^4^ cells per group). Stacked bar graph of the fraction of Tc cells in each state and transition. (E) Heat map of Tc cell densities in Palantir plots for KP LucOS following PBS treatment (Ctrl, vaccine cohort) separated by distance from tumor boundary (distal: >50 µm from the tumor boundary; proximal: <50 µm from the tumor boundary; and inside tumor, n = 10^4^ cells). (F) Heat map of Tc densities in Palantir projections for KP LucOS mice following Ctrl or ICB treatment separated by distance from tumor boundary as in E (n = 10^4^ cells per treatment). (G) Frequency of Tc cell states and transitions from tumor boundaries in ICB cohort. (H) Plot of the percent of Tc cells that are TCF1+ PD1+ in each Tc cell state in ICB cohort.

**Figure S6.**
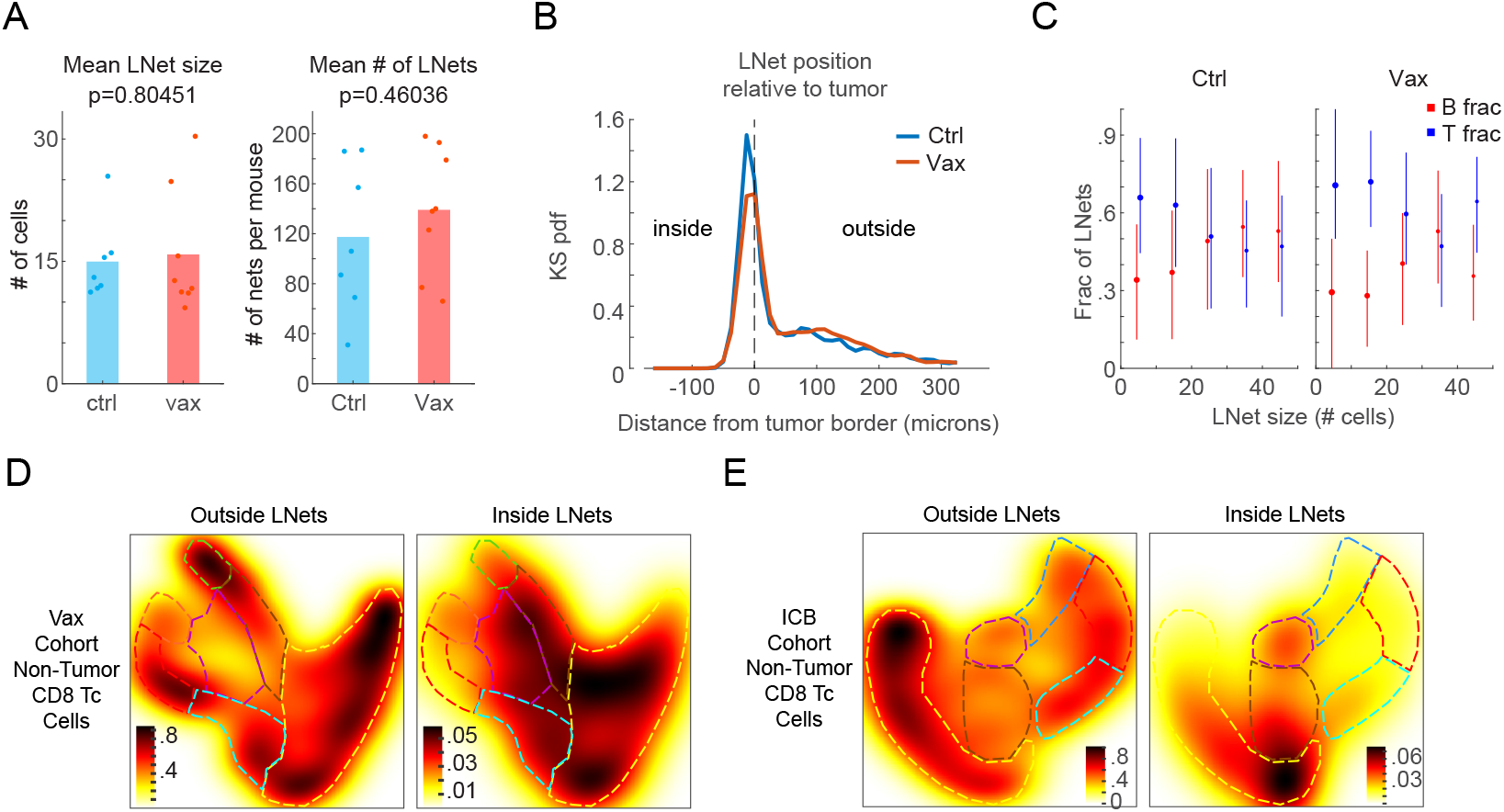
Characterization of lymphonets after vaccine and immune checkpoint blockade immunotherapies. (A) Plot of the number of cells in lymphonets and the average number of lymphonets in KP LucOS mice without treatment (Ctrl) or treated with vaccination (Vax) (n = 7 and 8 mice, bar = mean). (B) Kernel density probability density function of lymphonet spatial frequency relative to the tumor boundary for Ctrl (n = 7) and Vax (n = 8) mice. (C) Plot of the fraction of lymphonets comprised of T and B cells as a function of lymphonet size in Ctrl and Vax mice (mean +/- 25^th^ percentile). (D-E) Heat map of cell densities of non-tumor Tc cells present outside and inside lymphonets in Palantir projections for Vax (D, n = 14,480 and 978 cells) and ICB-treated (E, n = 13,948 and 735 cells) cohorts.

**Table S1:**
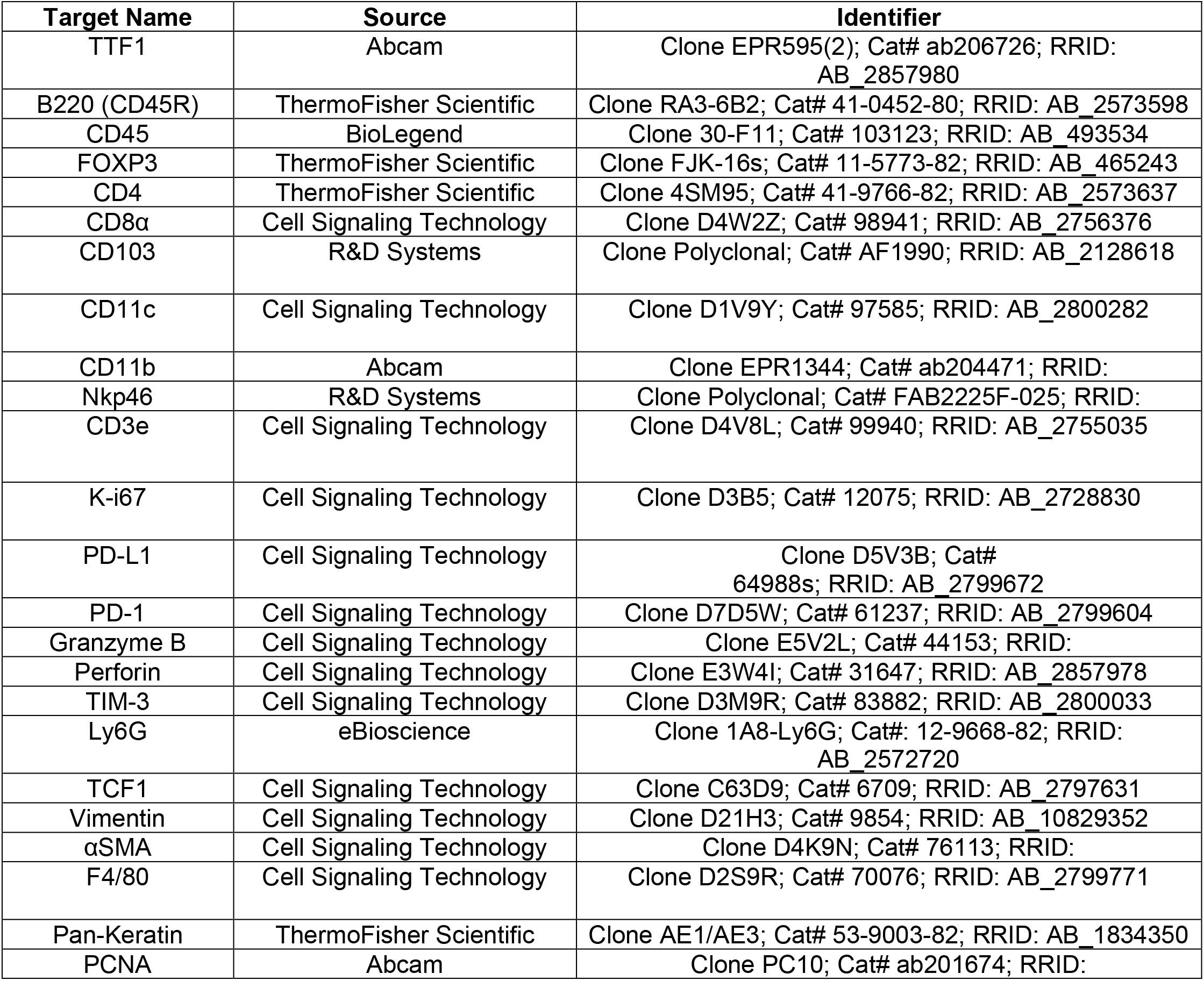
CyCIF Antibody Information

**Table S2:**
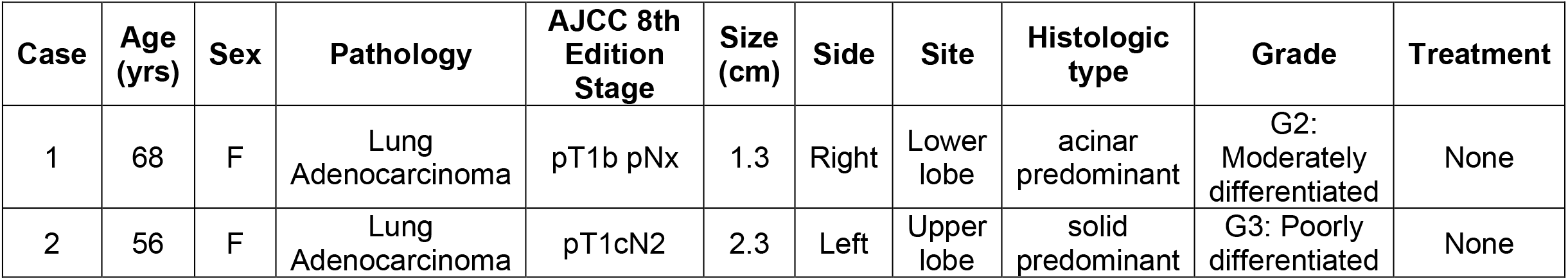
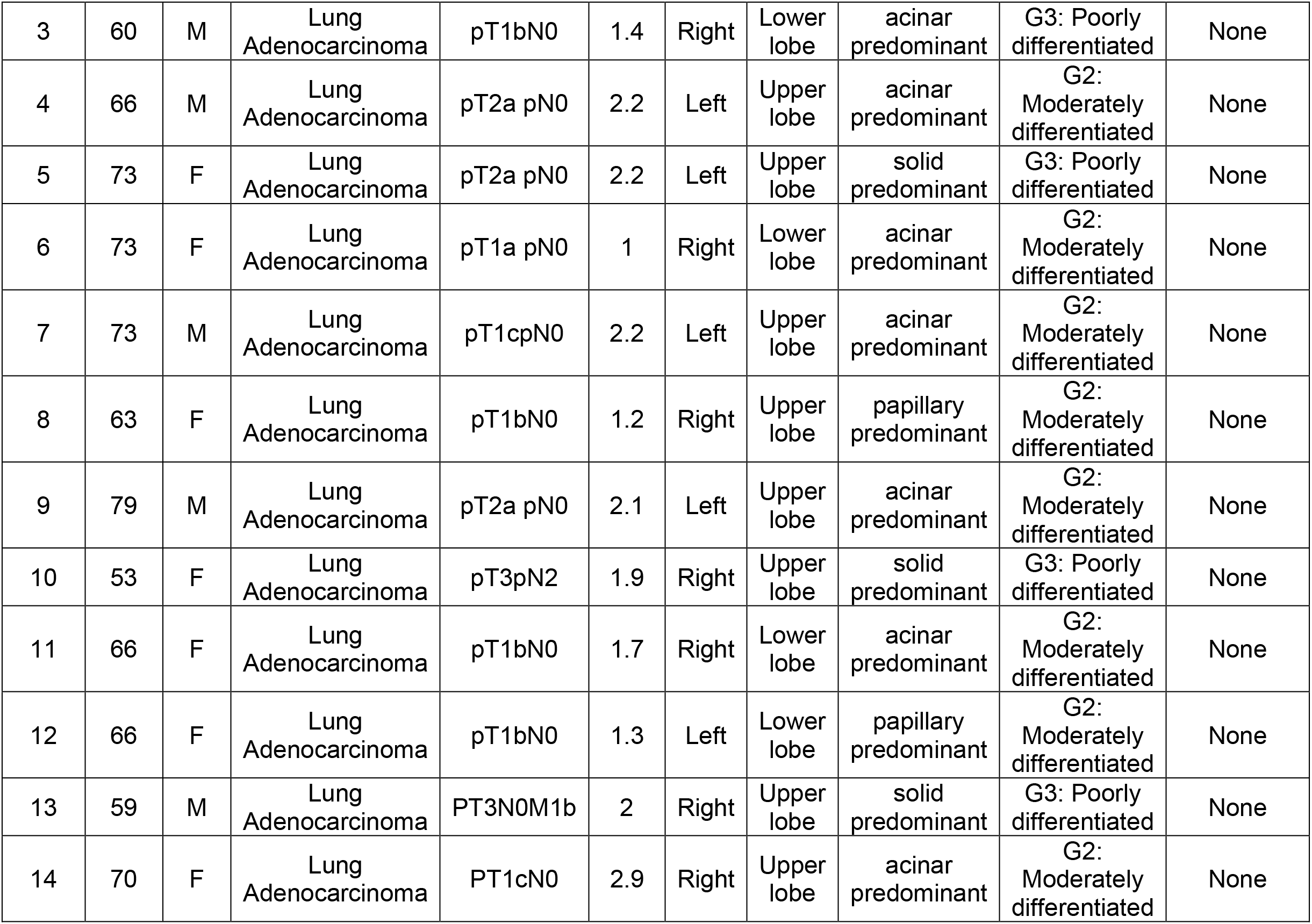
Human samples information

**Table S3:** Lymphonets and Tertiary Structures in Human Lung Adenocarcinoma

| CASE | Lymphonet number (by size) |  |  |  |  | TLS |
| --- | --- | --- | --- | --- | --- | --- |
|  | <10 cells | 11-30 | 31-64 | 65-1000 | >1000 |  |
| 1 | 1944 | 1009 | 151 | 133 | 18 | n/a |
| 2 | 1317 | 525 | 83 | 64 | 12 | 12 |
| 3 | 583 | 215 | 41 | 34 | 3 | 5 |
| 4 | 3052 | 1256 | 132 | 80 | 2 | 7 |
| 5 | 3679 | 1505 | 240 | 175 | 3 | 12 |
| 6 | 1487 | 658 | 115 | 68 | 1 | 4 |
| 7 | 1328 | 563 | 124 | 97 | 0 | 1 |
| 8 | 1581 | 773 | 158 | 154 | 8 | 36 |
| 9 | 2844 | 1347 | 214 | 121 | 12 | 18 |
| 10 | 1385 | 771 | 193 | 117 | 4 | 9 |
| 11 | 1666 | 946 | 187 | 118 | 10 | 12 |
| 12 | 832 | 379 | 74 | 43 | 3 | 8 |
| 13 | 2629 | 1420 | 253 | 114 | 6 | 13 |
| 14 | 859 | 503 | 103 | 94 | 6 | 66 |

**Table S4:** Mouse Experiment Information

| Expt # | Viral Construct | Treatment | # mice | Imaging modality |
| --- | --- | --- | --- | --- |
| 1 | Cre | none | 5 | CyCIF |
| 1 | LucOS | none | 5 | CyCIF |
| 1 | Cre+Cxcl10a | none | 5 | CyCIF |
| 2 | Cre | none | 5 | RNA Scope™ |
| 2 | LucOS | none | 5 | RNA Scope™ |
| 2 | Cre+Cxcl10a | none | 5 | RNA Scope™ |
| 3 | LucOS | PBS (control) | 6 | CyCIF |
| 3 | LucOS | ICB | 6 | CyCIF |
| 4 | LucOS | PBS (control) | 7 | CyCIF |
| 4 | LucOS | Vax | 8 | CyCIF |

